# Interaction of an α-synuclein epitope with HLA-DRB1*15:01 triggers enteric features in mice reminiscent of prodromal Parkinson’s disease

**DOI:** 10.1101/2022.02.03.479014

**Authors:** Francesca Garretti, Connor Monahan, Nicholas Sloan, Jamie Bergen, Sanjid Shahriar, Seon Woo Kim, Alessandro Sette, Tyler Cutforth, Ellen Kanter, Dritan Agalliu, David Sulzer

## Abstract

Enteric symptoms, including constipation, are hallmarks of prodromal Parkinson’s disease (PD) that can appear decades before the onset of motor symptoms and diagnosis. PD patients possess circulating T cells that recognize specific α-synuclein-(α-syn)-derived epitopes. One epitope, α-syn_32-46_, binds with strong affinity to the HLA-DRB1*15:01 allele implicated in autoimmune diseases. We report that α-syn_32-46_ immunization in a mouse expressing HLA-DRB1*15:01 triggers intestinal inflammation leading to loss of enteric neurons, damage of enteric dopaminergic neurons, constipation and weight loss. α-Syn_32-46_ immunization activates innate and adaptive immune gene signatures in the gut and induces changes in CD4^+^ TH1/ TH17 transcriptome that resemble tissue resident memory cells found in mucosal barriers during inflammation. Depletion of CD4^+^, but not CD8^+^, T cells partially rescues enteric neurodegeneration. Therefore, interaction of α-syn_32-46_ and HLA-DRB1*15:0 is critical for gut inflammation and CD4^+^ T cell-mediated loss of enteric neurons in humanized mice, suggesting potential mechanisms of prodromal enteric PD.

**HIGHLIGHTS AND eTOC Blurb:**

1. α-syn_32-46_ immunization of an HLA-DRB1*15:01 mouse triggers weight loss and constipation.
2. α-syn_32-46_ immunizations induce gut inflammation, loss of enteric neurons and damage to dopaminergic neurons.
3. α-syn_32-46_ immunization induces innate and adaptive immune responses in the gut.
4. Depletion of CD4^+^, but not CD8^+^, T cells partially rescues enteric neural loss.
5. An interaction between α-syn_32-46_ and HLA-DRB1*15:01 is critical for this model of prodromal PD.

Parkinson’s disease (PD) patients exhibit elevated number of circulating T cells that recognize α-synuclein-(α-syn)- epitopes, particularly during early disease stages. One epitope, α-syn_32-46_, interacts with the HLA-DRB1*15:01; however, its role in PD pathogenesis remains unknown. Garretti *et al*. show that α-syn_32-46_ immunization of a mouse expressing HLA-DRB1*15:01 triggers intestinal inflammation, a loss of enteric neurons, constipation and weight loss, suggesting a critical role for α-syn autoimmunity in HLA-DRB1*15:01 carriers in prodromal PD.

## INTRODUCTION

Parkinson’s disease (PD) is a neurodegenerative disorder characterized by prominent motor symptoms resulting from the loss of central nervous system (CNS) dopaminergic neurons located in the substantia nigra (SN) (Fahn and Sulzer, 2004). However, pathogenic steps in PD begin long before diagnosis, and the death of these neurons is suspected to occur at relatively late stages of disease pathogenesis. In contrast, enteric nervous system (ENS) symptoms are common during an earlier prodromal stage of the disease (Heinzel et al., 2019). Constipation presents in approximately 70% of PD patients (Fasano et al., 2015), is three-fold more prevalent in PD patients than healthy controls (Drossman, 2006), and may occur as early as twenty years prior to the onset of motor symptoms (Khoo et al., 2013). Although epidemiological studies have demonstrated a high prevalence of constipation in PD patients [reviewed in (Fasano et al., 2015)], and postmortem studies indicate that gut pathology is present before the onset of motor symptoms (Braak et al., 2003; Stokholm et al., 2016), the mechanisms underlying the enteric pathophysiology remain unclear.

Braak and colleagues have hypothesized that α-synuclein (α-syn) -mediated pathology begins in the ENS and subsequently ascends rostrally into the brain via the vagus nerve (Braak et al., 2003). This conjecture is consistent with reports that truncal vagotomies decrease the risk for PD (Liu et al., 2017; Svensson et al., 2015; Tysnes et al., 2015). Epidemiological studies support a role for gut inflammation in early stages of PD. The incidence of PD among patients with inflammatory bowel disease (IBD) is 28% higher than controls. Furthermore, IBD patients who receive anti-TNFα therapy exhibit a 78-100% reduction in PD incidence compared to those who do not receive such therapy (Park et al.; Peter et al., 2018). Multiple observations in animal models are also consistent with the hypothesis that gut inflammation may be involved in the early stages of PD-like pathogenesis [reviewed in (Metzger and Emborg, 2019)].

A possible role for peripheral inflammation in PD pathogenesis is suggested by findings that increased levels of pro-inflammatory cytokines (e.g., TNFα, IL-1β, IL-6 and IFNγ) are found in PD patients and correlate negatively with disease duration (Cossais et al., 2021; Devos et al., 2013; Pochard et al., 2018; Qin et al., 2016). PD patients also possess circulating T cells, mostly CD4^+^ subtypes, that recognize specific α-syn-derived neo-epitopes (Sulzer et al., 2017). These cells are present at particularly high levels during the first decade following diagnosis and likely during the prodromal phase prior to the onset of motor symptoms and clinical diagnosis (Lindestam Arlehamn et al., 2020). Thus, α-syn neo-antigen reactivity may be relevant to the prodromal and early stages of PD [reviewed in (Garretti et al., 2019)]. However, the role of α-syn neo-antigen reactivity in disease etiology remains unknown.

Previously, we reported that most α-syn-responsive CD4^+^ T cells isolated from PD patients respond to epitopes derived from two regions of the protein (Sulzer et al., 2017). One region requires the presence of phosphorylated amino acid serine 129 (Sulzer et al., 2017), a post-translational modification present at high levels in Lewy bodies (Fujiwara et al., 2002). However, these C-terminal epitopes bind weakly to several human leukocyte antigen (HLA) alleles, indicating that they are “unrestricted”. The other major α-syn epitope identified in PD patients is the peptide α-syn_32-46_, which lies in proximity to several rare α-syn mutations that cause familial PD (Appel-Cresswell et al., 2013; Ki et al., 2007; Kruger et al.; Lesage et al., 2013; Liu et al., 2021; Pasanen et al., 2014; Polymeropoulos et al., 1997; Yoshino et al., 2017; Zarranz et al., 2004). In contrast to the C-terminal epitopes, the α-syn_32-46_ epitope is highly restricted to a specific HLA haplotype [B*07:02, C*07:02, DRB5*01, DRB1*15:01, DQA1*01:02, DQB1*06:02], expressed by approximately one third of the population (Sulzer et al., 2017). Importantly, there is a very strong binding affinity (K_D_=2.8 nM) of the peptide to HLA-DRB1*15:01 *in vitro* (Sulzer et al., 2017). Independent genome-wide association studies (GWAS) have also confirmed that this haplotype is associated with PD (Hamza et al., 2010; Hill-Burns et al., 2011; Kannarkat et al., 2015).

It is unknown how the interaction between this α-syn epitope (α-syn_32-46_) and HLA allele (DRB1*15:01) may trigger early features of PD. Here, we report that α-syn_32-46_ peptide immunizations of a mouse strain lacking the mouse MHC-II alleles and expressing the human HLA-DRB1*15:01 allele trigger intestinal inflammation and loss of enteric neurons in the submucosal plexus of the small intestine leading to constipation and weight loss; however, there were no detectable effects in the CNS. Bulk RNA sequencing revealed that α-syn_32-46_ immunizations induced transcriptome changes in the gut related to innate and adaptive immune responses and interferon signaling. Single cell RNA sequencing (scRNAseq) of immune cells showed altered gene signatures in CD4^+^ T_H_1 and T_H_17 lamina propria lymphocytes of CFA/α-syn_32-46_ -immunized mice. These cells adopted a transcriptome signature characteristic of antigen-experienced tissue resident memory cells found in mucosal barriers during infection and chronic inflammation. These enteric pathologies and symptoms were not observed in wild-type mice immunized with the same peptide, indicating a specificity of the immune response to a specific MHC-II genotype. Depletion of CD4^+^, but not CD8^+^, T cells partially rescued enteric neurodegeneration. Thus, interactions of α-syn_32-46_ peptide with the HLA-DRB1*15:01 allele are critical for the induction of enteric features resembling those found in prodromal PD, and suggest that additional hits (e.g., blockade of intracellular protein turnover and/or α-syn aggregation) may be required for development of CNS symptoms.

## RESULTS

### α-Syn_32-46_ immunizations cause illness marked by severe weight loss and constipation in HLA-DRB1*15:01, but not wild-type, mice

Previously, we have shown that CD4^+^ T cells isolated from PD patients carrying the HLA-DRB1*15:01 allele confer a strong immune response to the α-syn_32-46_ region of the protein (Sulzer et al., 2017). To test if anti-α-syn_32-46_ immune responses identified in PD patients can produce features of PD pathology in mice, we performed active immunizations by adapting a protocol similar to the myelin oligodendrocyt glycoprotein (MOG_35-55_) experimental autoimmune encephalomyelitis (EAE) mouse model for human multiple sclerosis (Lengfeld et al., 2017; Lutz et al., 2017). Mice were immunized with either PBS with CFA (complete Freund’s adjuvant), or α-syn_32-46_ with CFA (referred to as CFA and CFA/α-syn_32-46_ immunization, respectively). As in EAE, *Bordetella pertussis* (*B. pertussis*) toxin was administered intravenously at 14 and 16 days post-initial immunization (DPI) with the boost to transiently open the blood-brain barrier in order to permit immune cell infiltration into the CNS (**Figure 1A**). In addition, mice were injected with PBS alone to control for the effects of CFA administration (referred to herein as PBS). These immunizations were performed either in C57BL/6J [referred to as wild-type (WT)] mice, or a humanized mouse strain lacking the mouse MHC-II alleles and expressing the human HLA-DRB1*15:01 allele [referred to as the HLA mice (Finn et al., 2004; Khare et al., 2005)] (**Figure 1A**). We confirmed that splenocytes from HLA-DRB1*15:01 mice expressed HLA-DR protein, and that its expression was increased when these cells were exposed to lipopolysaccharide (LPS) (**Figure S1A-G**). HLA-DR protein was not detected in WT splenocytes, as expected (**Figure S1H, I**). In addition, we confirmed that HLA mice expressed similar proportions of CD4^+^ and CD8^+^ T cells to WT mice (**Figure S1J-L**).

**Figure 1.**
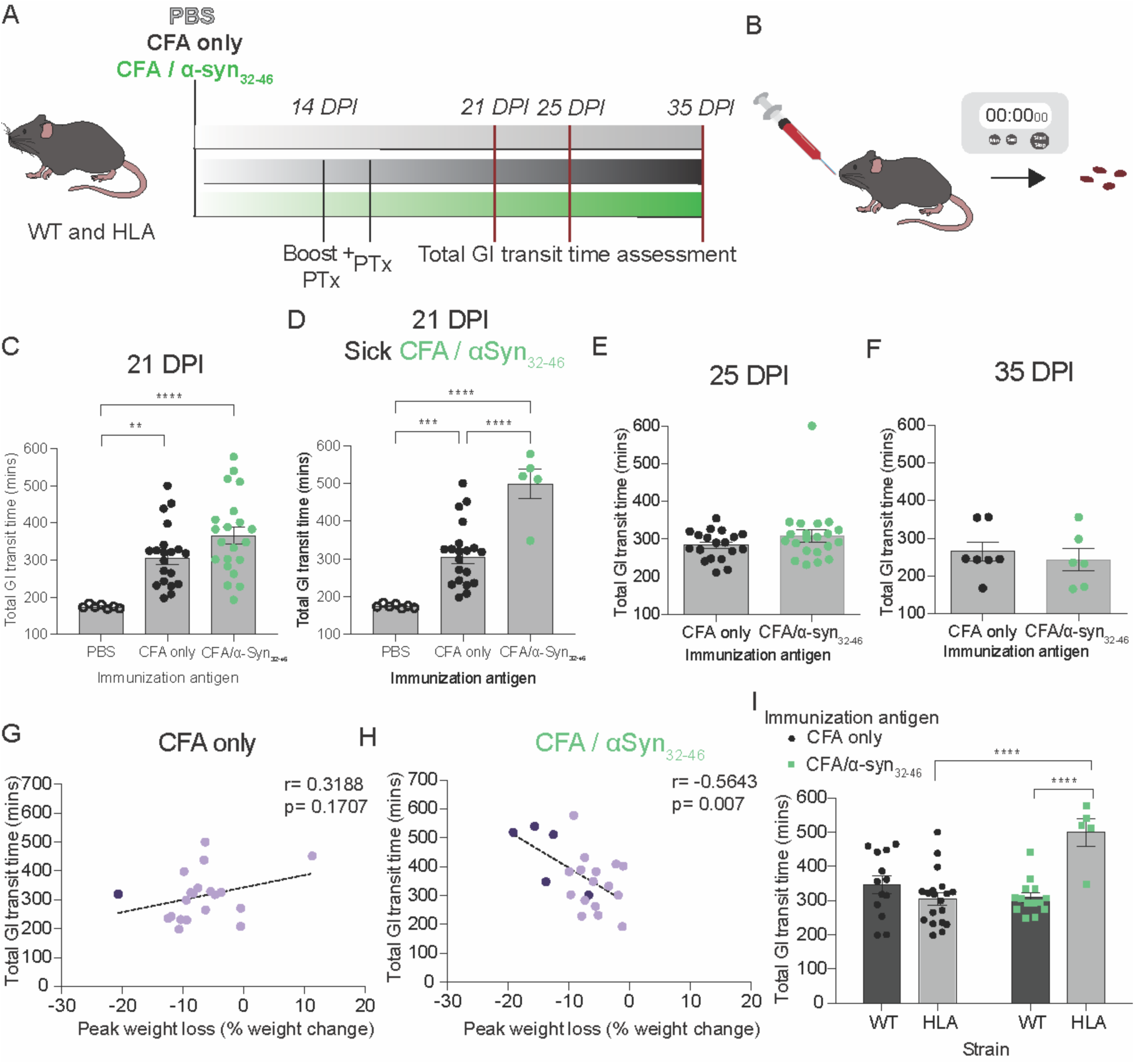
α-Syn_32-46_ immunizations induce weight loss in HLA DRB1*15:01, but not wild-type, mice. **(A)** A schematic diagram of the experimental design. Wild-type (WT) and HLA mice were immunized with PBS, CFA only, or CFA/ α-syn_32-46_ peptide and received a boost after two weeks. At 14 and 16 DPI, an intravenous injection of *B. pertussis* toxin (Ptx) was administered. Mice were monitored daily for signs of illness and change in weight. **(B)** Pie charts show the fraction of WT (top) - and HLA (bottom) - immunized mice that were healthy (light purple), became ill (dark purple) or died (gray). (**C, D**) Graphs of percentage weight change from the initial weight (0 DPI) for WT **(C)**, and HLA **(D)** mice immunized with either PBS (grey triangles), CFA only (black circles), or CFA/α-syn_32-46_ peptide (green triangles). The red dashed lines indicate the time when mice received an immunization boost and PTx injections at 14 and 16 DPI, respectively. **(E)** Graph depicting the percentage of weight loss for CFA only and sick CFA/α-syn_32-46_ HLA mice. The data in (**E**) were analyzed using a mixed-effect ANOVA for repeated measurements followed by Bonferroni post-hoc correction; * p<0.05. **WT**: CFA only (n= 23 mice), CFA/ α-syn_32-46_ peptide (n=25 mice). **HLA**: PBS (n=16 mice), CFA only (n=34 mice), CFA/ α-syn_32-46_ peptide (n=35 mice).

Mice were monitored daily for overt signs of malaise or weight changes (**Figure 1B-E**). Mice were categorized as “sick” if they lost more than 12% of the initial body weight and exhibited hunched posture with ungroomed and “ruffled” fur (**Figure 1B**). Following immunizations, very few WT mice became sick regardless of the antigen used for immunization (0% of CFA; 4% CFA/α-syn_32-46_; **Figure 1B, C**). In contrast, while CFA alone produced no illness in HLA mice, 25% of HLA mice immunized with CFA/α-syn_32-46_ became ill or died (**Figure 1B, D**). The “sick” CFA/α-syn_32-46_-immunized HLA mice exhibited severe weight loss starting at 16 DPI that peaked at 22-24 DPI, with a mean weight change of −15.68 % at 22 DPI (p= 0.0272; **Figure 1E**). The weight effects seen with CFA/α-syn_32-46_ immunization were specific to HLA mice and were absent in WT mice (**Figure 1C**). This phenotype was transient: sick HLA-immunized mice restored their initial weight between 24 - 29 DPI, with no further weight gain (**Figure 1E**).

As constipation is often an early symptom associated with PD that occurs as early as 20 years prior to the onset of motor symptoms (Khoo et al., 2013), we assessed total gastrointestinal transit time in either PBS, CFA, or CFA/α-syn_32-46_-immunized mice by measuring the time required for a non-absorbable red dye to be excreted with feces after oral administration (**Figure 2A, B**). We selected three times points after immunization (21, 25 and 35 DPI) to correlate this phenotype with the weight loss (**Figure 1E)**. Immunizations with CFA had a significant effect on total GI transit time (PBS vs CFA, p= 0.0029; PBS vs CFA/α-syn_32-46_ p<0.0001; **Figure 2C, D**). While the GI transit time between all CFA/α-syn_32-46_- and CFA-immunized HLA mice at 21 DPI was not significantly different (p= 0.1473; **Figure 2C**), the sick CFA/α-syn_32-46_ HLA mice displayed significantly longer GI transit time compared to both PBS and CFA mice (p< 0.0001; **Figure 2D**). The increase in GI transit time, however, recovered by 25 and 35 DPI (**Figure 2E, F**), mirroring the recovery from weight loss (**Figure 1E**). Interestingly, all CFA/α-syn_32-46_ HLA mice showed a significant negative correlation between the peak weight loss and GI transit time (r= −0.5643, p= 0.0077; **Figure 2H**), whereas this was not seen with CFA-immunized HLA mice (peak weight loss and GI transit time; r= 0.3188, p= 0.1707; **Figure 2G**). In contrast, all immunized WT mice exhibited unaltered total GI transit times, regardless of the immunization antigen, indicating a specificity of the immune response (**Figure 2I**). Thus, the combination of α-syn_32-46_ peptide and HLA-DRB1*15:01 allele expression induces acute gastrointestinal illness (weight loss and constipation) between 22-24 DPI.

**Figure 2.**
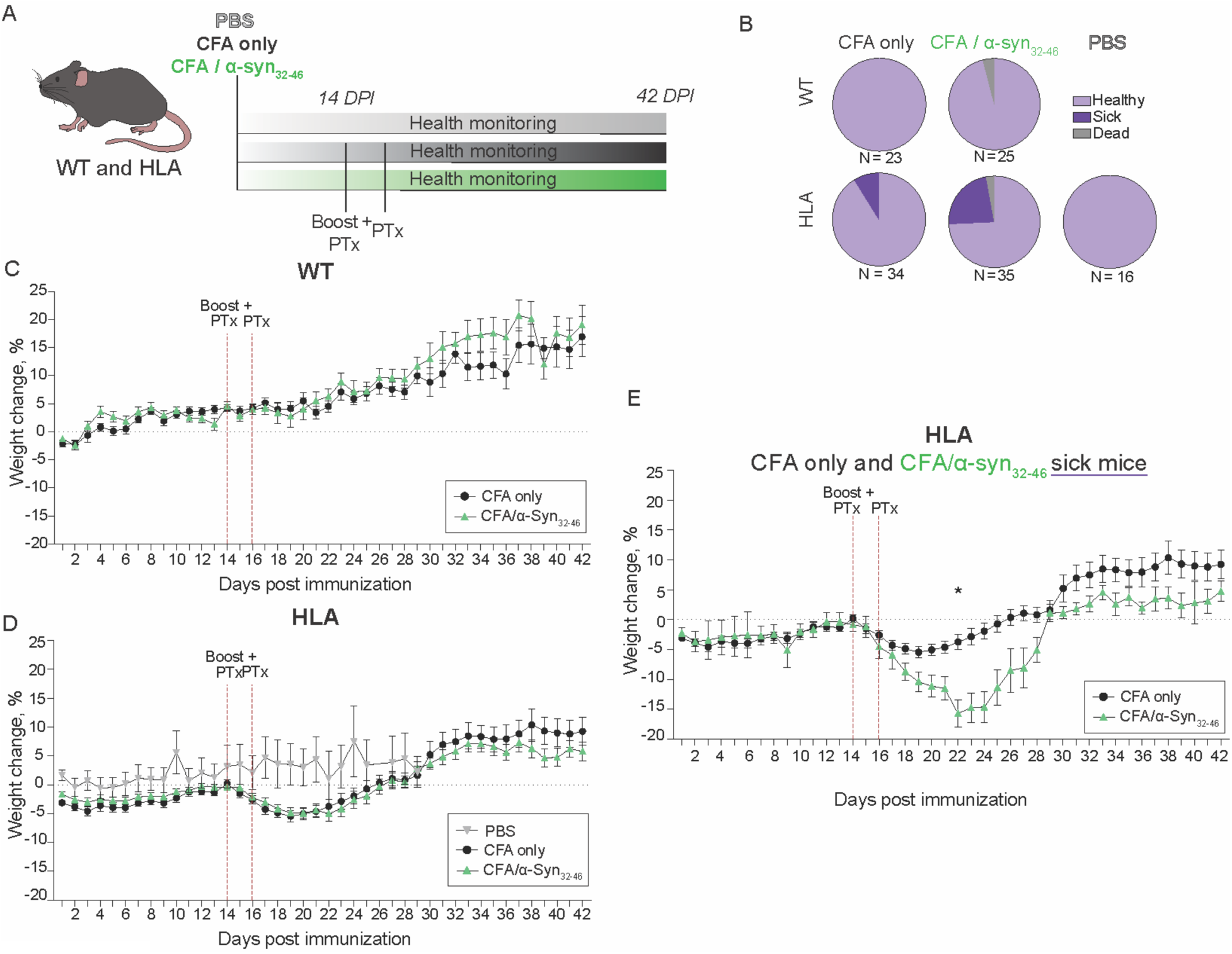
Total GI transit time is delayed in CFA/α-syn_32-46_ -immunized HLA mice. **(A, B)** Schematic diagrams of the experimental design for immunization and gastrointestinal (GI) transit time. Mice were gavaged with a non-absorbable carmine red mixture. The time from the oral gavage to the release of a red fecal pellet was recorded as the total gastrointestinal (GI) transit time. The total GI time was measured at 21, 25, and 35 days post-immunization (DPI). (**C, D**) Dotted bar graphs depicting the total GI transit time at 21 DPI for either all **(C)**, or sick **(D)**, PBS (empty circles), CFA only (black) and CFA/α-syn_32-46_ (green)-immunized HLA mice. (**E, F**) Dotted bar graphs depicting the total GI transit time of CFA only (black) and CFA/α-syn_32-46_ (green)-immunized HLA mice at 25 **(E)** and 35 **(F)** DPI. (**G, H**) Correlation graphs between peak weight loss and total GI transit time of CFA only **(G)**, and CFA/α-syn_32-46_ **(H)** - immunized HLA mice. Dark purple circles represent sick and light purple circles represent healthy mice. **(I)** Bar graphs of the total GI transit time of WT (dark bars) and HLA (light bars) immunized with CFA only (black circles) or CFA/α-syn_32-46_ (green circles). Only sick CFA/α-syn_32-46_-immunized HLA mice are shown. Data were analyzed by Mann-Whitney test (**C, D**), Pearson correlation (**G, H**) and two-way ANOVA (**I**); ** p<0.01, *** p<0.001, **** p<0.0001. Dotted bar graphs represent the mean and SEM. Each symbol represents data from one animal. **WT**: CFA only n=13 mice, CFA/α-syn_32-46_ n=14 mice. **HLA**: PBS n=7 mice, CFA only n=20 mice, CFA/α-syn_32-46_ n=21 mice. WT: 2 independent experiments, HLA: 5 independent experiments.

### α-Syn_32-46_-immunized HLA mice exhibit loss of the enteric, but not central nervous system, neurons

The weight loss associated with constipation observed in CFA/α-syn_32-46_-immunized HLA mice suggests a pathological response targeting the gut. To examine the gut histopathology underlying these phenotypes, we prepared flat mounts of both the submucosal and myenteric plexuses of the ileum (small intestine) from immunized HLA mice at 28 DPI (**Figure 3A, B**), since this preparation provides an in-depth analysis of the ENS. We focused in particular on the submucosal plexus since it is directly exposed to circulating immune cells and has the highest concentration of dopaminergic neurons (Li et al., 2004). To investigate whether the increase in total GI transit time was associated with neuronal loss, we analyzed expression of ANNA1 as a marker for enteric neuronal nuclei, and tyrosine hydroxylase (TH) as a marker for catecholaminergic neurons, which in the ileum are mostly dopaminergic (Li et al., 2004).

**Figure 3.**
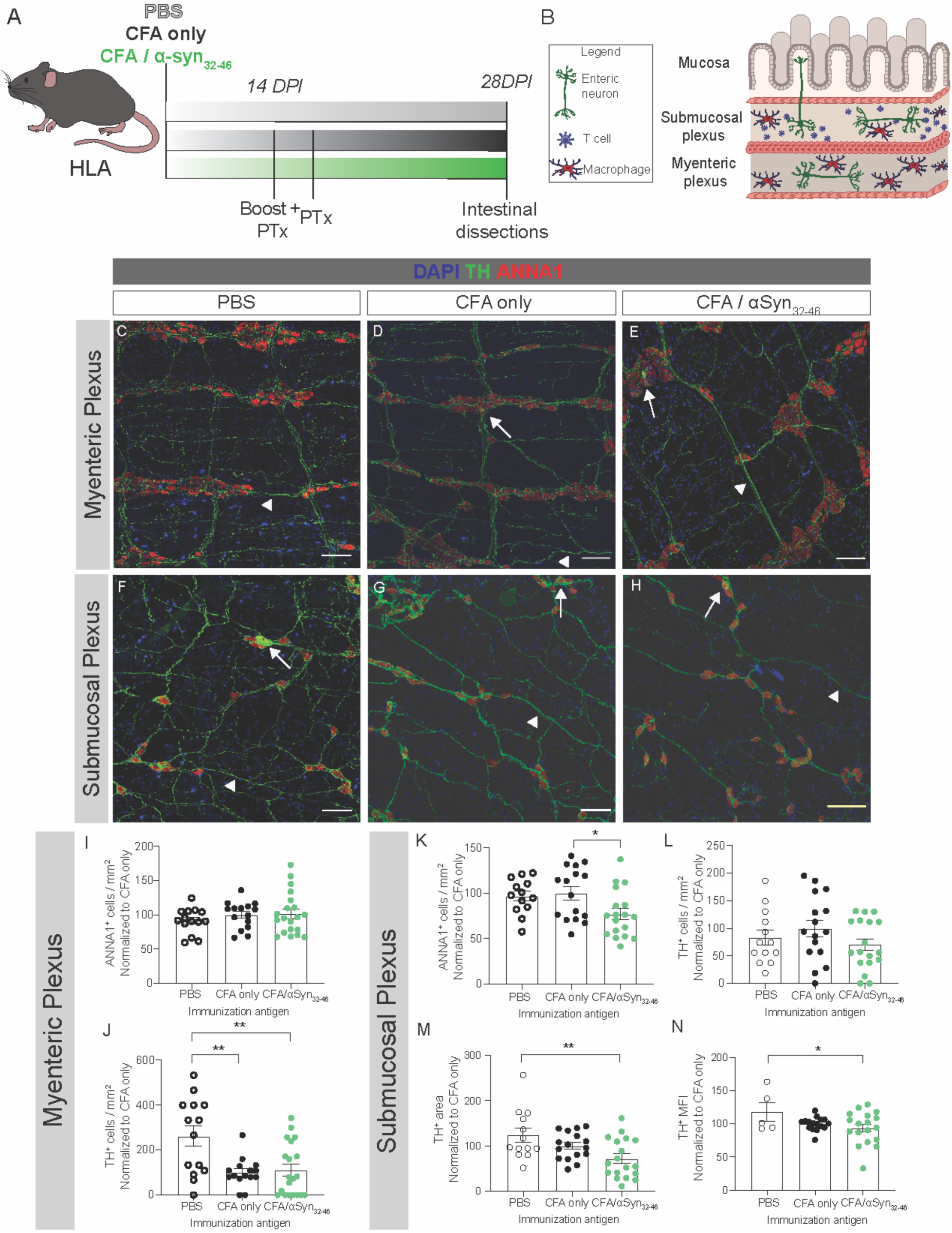
α-Syn_32-46_ immunizations induce enteric neuron loss in HLA DRB1*15:01 mice. **(A, B)** Schematic diagram of the experimental design and the myenteric and submucosal plexuses of the ileum used for the flat mount analysis. (**C-H**) Representative images of the myenteric **(C-E),** and submucosal (**F-H**) plexuses stained for DAPI (blue), TH (green), and ANNA1 (red). Thin white arrows indicate cell bodies of dopaminergic neurons; white arrowheads indicate dopaminergic neuronal processes. Scale bars = 75 μm. (**I-N**) Dotted bar graphs of ANNA1^+^ **(I)** and TH^+^ **(J)** neurons per mm^2^ in the myenteric plexus, ANNA1^+^ **(K),** or TH^+^ **(L)** neurons per mm^2^, area covered **(M)** and mean fluorescence intensity (MFI) **(N)** of the TH signal in the submucosal plexus of PBS (empty circles), CFA only (black circles) and CFA/α-syn_32-46_ (green circles)-immunized HLA mice. Dotted bar graphs show the mean and error bars the SEM. Data were analyzed by one-way ANOVA test. * p<0.05, ** p<0.01. PBS, n= 14; CFA only n=15 mice; CFA/α-syn_32-46_ n=20 (myenteric) and 18 (submucosal) HLA mice; data collected from 6 independent experiments.

In the myenteric plexus, there was a similar density of ANNA1^+^ neurons among PBS, CFA, and CFA/α-syn_32-46_ - immunized HLA mice (**Figure 3C-E, I**). The CFA immunization reduced the overall TH^+^ neuron density in the myenteric plexus (PBS vs CFA, p= 0.005; PBS vs CFA/α-syn_32-46,_ p=0.006; **Figure 3J**), consistent with adjuvant effects on total GI transit time (**Figure 2C, D**). In contrast, CFA/α-syn_32-46_-immunized HLA mice exhibited a significantly reduced ANNA-1^+^ neuronal density in the submucosal plexus compared to CFA-immunized HLA mice (p= 0.04, **Figure 3F-H, K**); however, there was no significant difference in ANNA-1^+^ neurons between PBS and CFA-immunized mice (**Figure 3K**). There was no significant difference in the number of TH^+^ neurons within the submucosal plexus in any group (**Figure 3L,** p= 0.260), which may be due to the variable density of TH^+^ neurons, despite the large areas of the ENS encompassed by the flat mount analysis. Importantly, the area of TH^+^ processes was significantly lower in CFA/α-syn_32-46_-immunized compared to PBS-injected HLA mice (**Figure 3M**, p=0.008), although there was no significant difference between CFA and CFA/α-syn_32-46_-immunized HLA mice (**Figure 3M**; p= 0.181). Similarly, the mean fluorescent intensity (MFI) of TH label within neuronal cell bodies and processes at 28 DPI was significantly lower in CFA/α-syn_32-46_ immunized than PBS HLA mice (**Figure 3N,** p= 0.017), although there was no significant difference between CFA and CFA/α-syn_32-46_-immunized HLA mice (**Figure 3N**; p= 0.171). The effects of CFA/α-syn_32-46_-immunizations on enteric neuronal loss were specific to HLA mice, and not seen in WT mice (**Figure S2)**. Thus, α-syn_32-46_ immunizations induced loss of enteric neurons in the ileum only in the HLA-DRB1*15:01 strain.

The submucosal plexus is exposed to circulating immune cells including T lymphocytes (**Figure 3B**) (Yoo and Mazmanian, 2017). To determine if the enteric neuron loss observed in CFA/α-syn_32-46_ immunized HLA mice correlates with immune cell infiltration, we immunolabeled and quantified Iba1^+^/CD68^+^ macrophages in these layers (**Figure S3**). We found no significant difference in CD68^+^ macrophage density in either the myenteric (**Figure S3 A-C, G**), or the submucosal (**Figure S3 D-F, H**) plexuses of the HLA mice regardless of the immunizing condition. Thus, the loss of enteric neurons does not correlate with the macrophage number in CFA/α-syn_32-46_ -immunized HLA mice.

Next, we examined whether CFA/α-syn_32-46_-immunized HLA mice exhibiting weight loss and constipation had CNS pathology. α-Syn_32-46_ immunizations did not induce CD3^+^ T cell infiltration, microglia activation, macrophage infiltration, or loss of SN dopaminergic neurons in the brains of either WT or HLA mice, up to two months after immunization (70 DPI; **Figure S4A-O**). Consistently, neither WT nor HLA mice displayed motor behavior or learning deficits after α-syn_32-46_ immunizations (**Figure S4P-Y**). The lack of CNS lymphocyte infiltration, neuroinflammation and neurodegeneration coupled with no changes in either motor skills or learning behaviors indicate that the severe weight loss and constipation in α-syn_32-46_-immunized HLA mice are likely not related to changes in basal ganglia circuits. Therefore, the responses to α-syn_32-46_ immunizations were restricted to the ENS in HLA mice.

### Dopaminergic neuron damage in the gut corresponds to and persists after the weight loss in α-syn_32-46_ - immunized HLA mice

To uncover the chronology between weight loss, gut inflammation and ENS neuronal death in CFA/α-syn_32-46_-immunized HLA mice, we analyzed the number of ANNA1^+^ and TH^+^ neurons in the myenteric and submucosal plexuses at 18 and 42 DPI (**Figure 4A and B**), prior to and after the weight loss (**Figure 1E**). At 18 DPI, there was no difference in the number of ANNA1^+^ neurons, dopaminergic neurons or TH^+^ fluorescence signal in the submucosal plexus between CFA and CFA/α-syn_32-46_-immunized HLA mice (**Figure 4E, F, J-L**). At 42 DPI, the area and MFI of the TH^+^ signal were both significantly decreased in the submucosal plexus of CFA/a-syn_32-46_ compared to CFA-immunized HLA mice (p= 0.0135 and p= 0.0066, respectively; **Figure 4O, P, U, V**), with no significant difference in the number of either ANNA1^+^ or dopaminergic TH^+^ neuronal bodies (p= 0.226; **Figure 4O, P, T**). There were no changes in any of these parameters in the myenteric plexus at 18 or 42 DPI (**Figure 4C, D, G, H, M, N, Q, R**). There was also no significant difference in the number of macrophages in the myenteric and submucosal plexuses at 18 and 42 DPI between CFA and CFA/α-syn_32-46_-immunized HLA mice (**Figure S3I-T**), suggesting no relationship between macrophage number and ENS neuron loss. Overall, the time course analysis indicates that damage of TH^+^ dopamine neurons in the submucosal plexus of CFA/α-syn_32-46_-immunized HLA mice began prior to 28 DPI, and persisted by day 42 DPI, after the mice have recovered from the weight loss, independent of macrophage infiltration in the gut.

**Figure 4.**
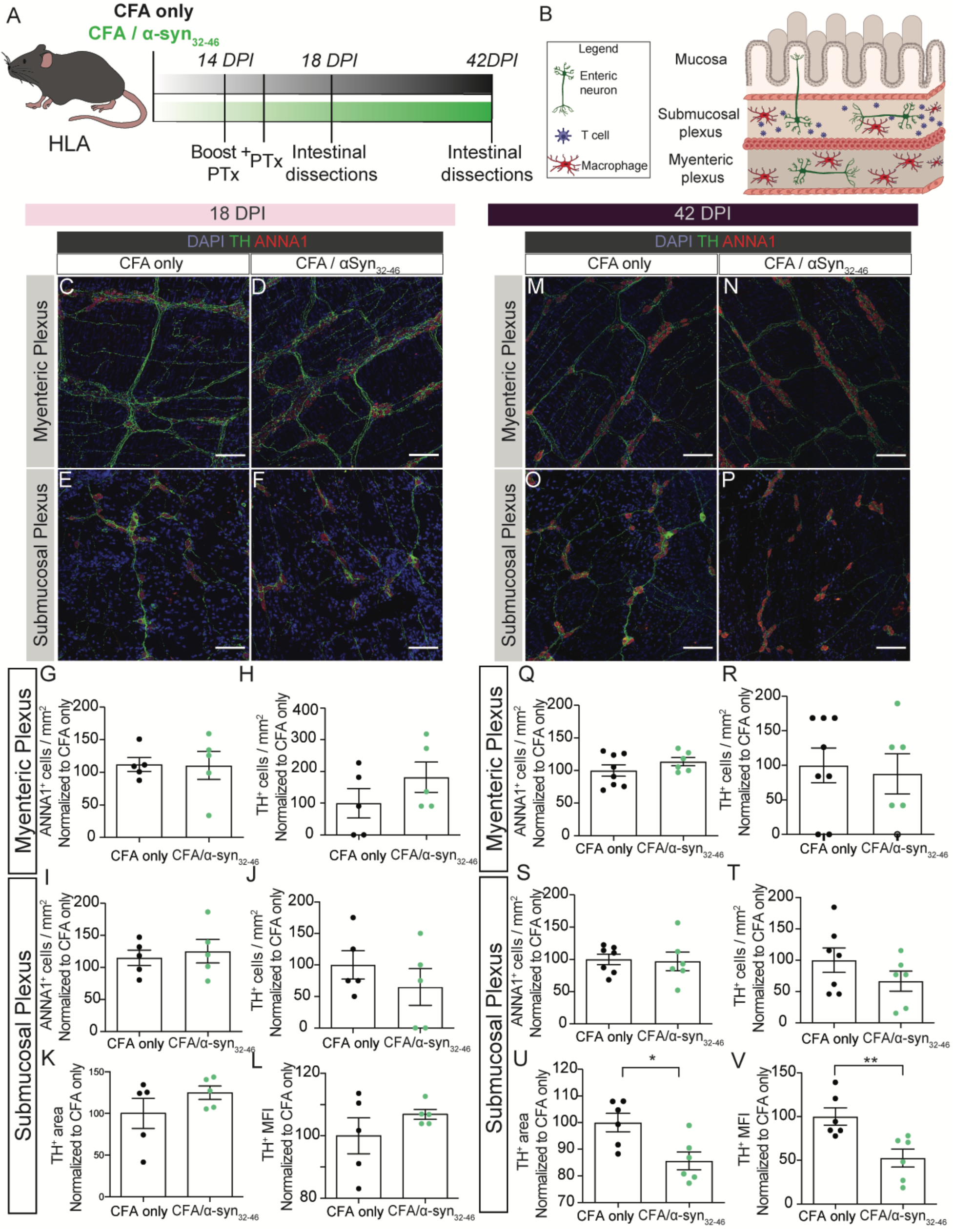
Neuronal loss begins after 18 days post-immunization in CFA/α-syn_32-46_ -immunized HLA DRB1*15:01 mice. **(A, B)** Schematic diagram of the experimental design (**A**) and the myenteric and submucosal plexuses of the ileum (**B**) used for analysis. HLA DRB1*15:01 mice were immunized with CFA only or CFA**/**α-syn_32-46_ and received a boost after two weeks. At 14 and 16 DPI, *B. pertussis* toxin (Ptx) was administered intravenously. At 18 and 42 DPI, the ileum was dissected out for analysis. **(C-F)** Representative images of the myenteric and submucosal plexuses at 18 DPI stained for DAPI (blue), TH (green), and ANNA1 (red). (**G, H**) Dotted bar graphs depicting the number of ANNA1^+^ **(G)** and TH^+^ **(H)** neurons in the myenteric plexuses of CFA only (black circles) and CFA/α-syn_32-46_ (green circles)-immunized mice at 18 DPI. (**I, J**) Dotted bar graphs of ANNA1^+^ **(I)** and TH^+^ cells **(J)** in the submucosal plexuses of CFA only (black circles) and CFA/α-syn_32-46_ (green circles)-immunized mice at 18 DPI. (**K, L**) Dotted bar graphs of the TH area **(K)** and mean fluorescence intensity (MFI) **(L)** of TH signal in the submucosal plexus of CFA only (black circles) and CFA/α-syn_32-46_ (green circles)-immunized mice at 18 DPI. **(M-P)** Representative images of the myenteric and submucosal plexuses at 42 DPI stained for DAPI (blue), TH (green), and ANNA1 (red). (**Q, R**) Dotted bar graphs of ANNA1^+^ **(Q)** and TH^+^ cells **(R)** in the myenteric plexus of CFA only (black circles) and CFA/α-syn_32-46_ (green circles)-immunized mice at 42 DPI. (**S-V**) Dotted bar graphs of ANNA1^+^ **(S)**, and TH^+^ cells **(T), TH** area **(U)** and mean fluorescence intensity (MFI) **(V)** of TH signal in the submucosal plexus of CFA only (black circles) and CFA/α-syn_32-46_ (green circles)-immunized mice at 42 DPI. Dotted bar graphs show the mean and error bars the SEM. Each symbol represents data collected from a mouse. Data were analyzed by a two-tailed Student t-test. * p<0.05, ** p<0.01. 18 DPI: CFA only n=5 mice, CFA/α-syn_32-46_ n=5 mice; 42 DPI: CFA only n=6-7 mice, CFA/α-syn_32-46_ n=6 mice (data were collected from one experiment). (**C-F; M-P**) Scale bars = 50μm.

### α-Syn_32-46_ immunization activates innate and adaptive immune gene signatures in the gut

To investigate the mechanisms underlying the gut inflammation in CFA/α-syn_32-46_ -immunized HLA mice, we performed bulk RNA sequencing and differential gene expression analysis of the ileum between CFA/α-syn_32-46_ and CFA-immunized HLA mice at 21 DPI, when mice showed the largest weight loss (**Figure 5A; Table S1**). Differential gene expression analysis identified 143 upregulated, and 54 downregulated genes in CFA/α-syn_32-46_ - immunized HLA mice (**Figure 5B**; p_adj_ < 0.05 and |log_2_fc| > 0.5; **Table S1B-D**). Gene ontology (GO) enrichment analysis using the Database for Annotation, Visualization and Integrated Discovery (DAVID) revealed significant GO terms for genes with similar functional characteristics upregulated in CFA/α-syn_32-46_ immunized HLA mice such as: a) the innate immune response (*e.g.*, GO: 0045087, FDR = 2.34 x 10^-25^); b) the response to interferon-β (GO: 0035456, FDR = 3.64 x 10^-15^) including interferon-stimulated genes (*Stat1/2*, *Oas2*, *Irf7*, and *Isg15*) that play multiple roles in defense against pathogens; and c) the response to cytokines (GO: 0034097, FDR = 3.54 x 10^-16^) including several chemokines (e.g., *Cxcl10* and *Cxcl13*) and *Vcam1*, a cell adhesion molecule involved in immune cell adhesion to blood vessels (**Figure 5C; Table S1E**). Additionally, genes involved in lymphocyte-mediated immunity (GO: 0002449, FDR = 0.00593) were also significantly enriched including antigen processing and presentation via MHC class I (*Tap1*, *H2.T24*, *H2.Q6*, *H2.T22*, *H2.T23*) (**Figure 5C; Table S1E**). Finally, humoral immune response was a significant GO term (GO: 0002455, FDR = 0.0495) with upregulation of genes encoding for the immunoglobulin heavy chain and complement cascade (**Figure 5C; Table S1E**). Thus CFA/α-syn_32-46_ immunizations upregulated innate and adaptive immune gene signatures in the ileum of HLA mice at peak weight loss and constipation.

**Figure 5.**
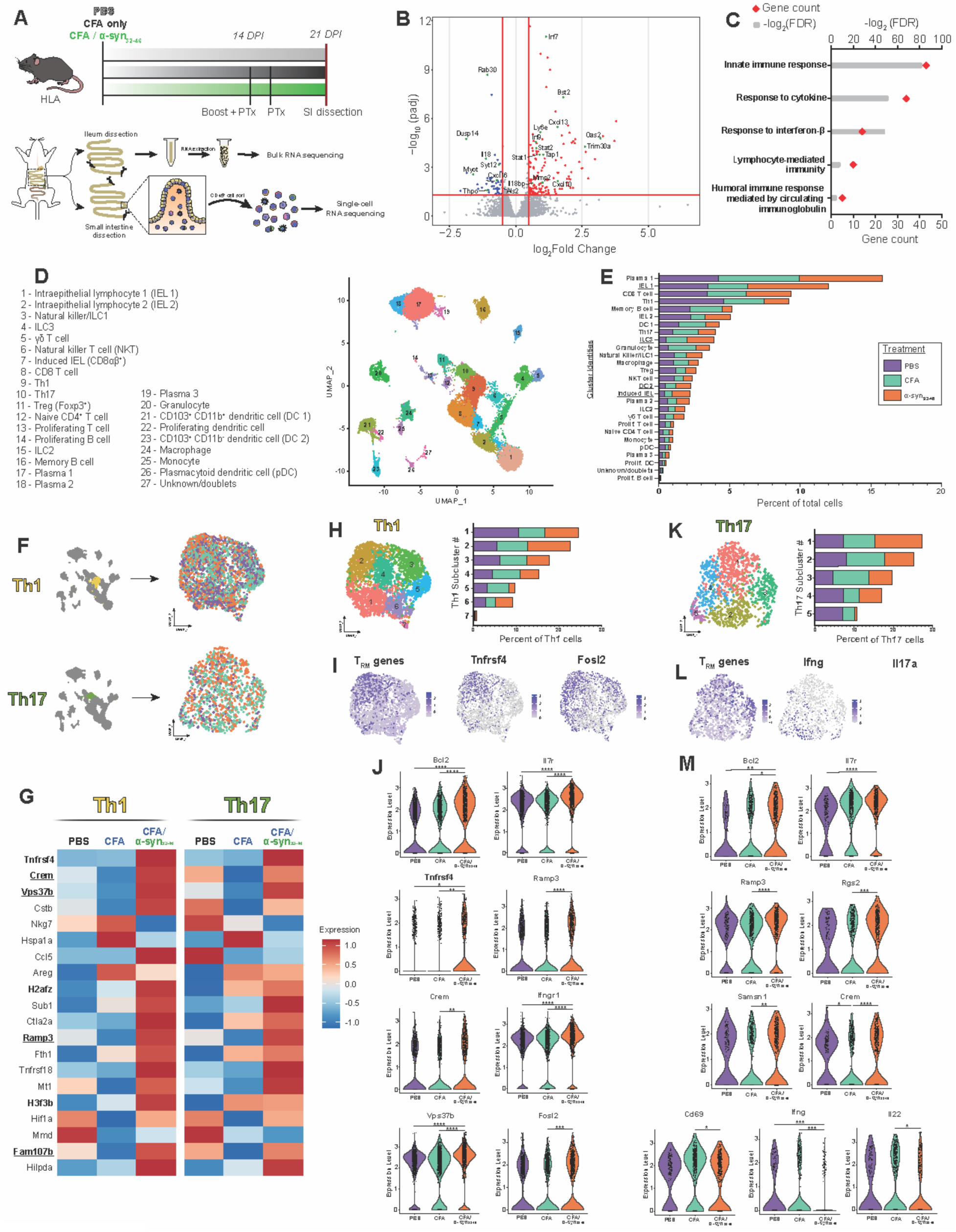
Gene signatures associated with the innate and adaptive immune responses and altered T_H_1/T_H_17 gene signatures are present in the small intestine of α-syn_32-46_-immunized HLA mice. **(A)** Schematic diagram of the experimental design for bulk and single-cell RNA seq (bulk RNA sequencing: CFA only n=4 mice, CFA/α-syn_32-46_ n=4 mice; single cell RNA sequencing: PBS n=2 mice, CFA n=3 mice, CFA/α-syn_32-46_ n=3 mice). **(B)** A volcano plot of differentially expressed genes from bulk RNA seq of the ileums of CFA/α-syn_32-46_ versus CFA only- immunized HLA mice at 21 DPI. Significantly down-regulated (blue) and up-regulated (red) genes are plotted by fold change on the x-axis and the adjusted p-value on the y-axis. Red lines indicate adjusted p values (p_adj_) ≤ 0.05 and |log_2_ fold change| ≥ 0.5 which were used to identify statistically significant, differentially expressed genes. Genes of interest are highlighted in green and labeled. **(C)** GO enrichment bar graph depicting five functional categories of upregulated genes from the ileum of α-syn_32-46_ immunized HLA mice at 21 DPI. Red diamonds indicate the number of genes represented by the GO term, and gray bars indicate -log_2_(FDR) values. GO terms are considered significant with a false discovery rate (FDR) < 0.05. **(D)** UMAP representation of single cell RNA sequencing (scRNAseq) data of CD45^+^ cells sorted from the small intestine of all three conditions (PBS, CFA, CFA/α-syn_32-46_). **(E)** Bar graph showing the proportion of sequenced CD45^+^ cells for each cluster of the UMAP (1-27 cell populations). Each bar shows the relative proportions of CD45^+^ cells for each condition [PBS (purple), CFA only (teal), CFA/α-syn_32-46_ (orange)] corrected for the number of cells sequenced from each condition. The underlined clusters are overrepresented (>45% of cluster) by cells isolated from CFA/α-syn_32-46_ HLA mice. **(F)** UMAP projections of T_H_1 and T_H_17 clusters isolated from the dataset [PBS (purple), CFA only (teal), CFA/α-syn_32-46_ (orange)]. **(G)** Heatmap of the average expression of 20 tissue-resident-memory (T_RM_)-associated genes in Th1 and Th17 cells, separated by condition. Bolded gene names are significantly upregulated in the Th1 population (p_adj_ < 0.05) from CFA/α-syn_32-46-_ compared to CFA only-immunized HLA mice. Underlined gene names are significantly upregulated in the T_H_17 population (p_adj_ < 0.05) in CFA/α-syn_32-46_ compared to CFA only-immunized HLA mice. **(H)** UMAP projection of the T_H_1 cell subset with a bar graph of the proportion of T_H_1 subclusters as a percent of total T_H_1 cells by condition. Each bar shows the relative proportions of cells from each condition [PBS (purple), CFA only (teal), CFA/α-syn_32-46_ (orange)] corrected for the number of total T_H_1 cells from each treatment. **(I)** Feature plots of T_RM_ genes from panel **G**, *Tnfrsf4*, and *Fosl2* in the T_H_1 cluster. **(J)** Violin plots of *Bcl2*, *Il7r*, *Tnfrsf4*, *Ramp3*, *Crem*, *Ifngr1*, *Vps37b*, and *Fosl2* gene expression in the T_H_1 cluster, separated by condition. Statistical analyses were done by Wilcoxon Rank Sum test, followed by Bonferroni correction for multiple comparisons. **(K)** UMAP projection of the T_H_17 cell subset with a bar graph of the proportion of T_H_17 subcluster as a percent of total T_H_17 cells by condition. Each bar shows the relative proportions of cells from condition [PBS (purple), CFA only (teal), CFA/α-syn_32-46_ (orange)] corrected for the number of cells from each treatment within the T_H_17 cluster. **(L)** Feature plots of T_RM_ genes (**from** Fig. 5G), *Ifng*, and *Il17a* in the T_H_17 cluster. **(M)** Violin plots of *Bcl2*, *Il7r*, *Ramp3*, *Rgs2*, *Samsn1*, *Crem*, *Cd69*, *Ifng*, and *Il22* gene expression in the T_H_17 cluster, separated by immunization treatment. For statistical significance: * p_adj_ < 0.05, ** p_adj_ < 0.01, *** p_adj_ < 0.001, **** p_adj_ < 0.0001.

### α-Syn_32-46_-specific T cells are present in the lymphatic organs of WT, but not HLA-DRB1*15:01 mice, after immunization

Since CFA/α-syn_32-46_ immunizations upregulate innate and adaptive immune gene signatures in the ileum of HLA mice, we examined the antigen-specific adaptive immune response. We isolated primary cells from either spleen or cervical and axillary lymph nodes of WT and HLA mice at 28 DPI after immunizations with either CFA or CFA/α-syn_32-46_ (**Figure S5A**). Cells were stimulated *in vitro* for 48 hours with either the peptide, PBS or Concanavalin A (ConA), as respective negative and positive controls. We then measured IFNγ and IL-17A levels, secreted by T_H_1 and T_H_17 lymphocytes, which are also elevated in the blood of PD patients (Sommer et al., 2018; Yang et al., 2014). α-Syn_32-46_ and ConA induced IFNγ and IL-17A secretion from WT splenocytes and lymph cells, but had no effect on cells from CFA and CFA/α-syn_32-46_-immunized HLA mice (p= 0.0691; **Figure S5B, C, G, H**; data not shown). The low levels of IFNγ in CFA/α-syn_32-46_ HLA splenocytes were released primarily by CD4^-^ CD8^-^ cells (p<0.0001), in contrast to the response of WT splenocytes after *in vitro* re-stimulation with α-syn_32-46_, which was largely driven by CD4^+^ T cells (p= 0.0160; **Figure S5D-F**). Thus, CD4^+^ T cells secreting IFNγ and IL-17A were not present in the spleens or lymph nodes in CFA/α-syn_32-46_-immunized HLA mice at 28 DPI, a response consistent with either release from these organs, or loss due to cell death after *in vitro* re-stimulation.

To determine if α-syn_32-46_ immunizations induced a humoral immune response in HLA mice, we assessed the levels of antibodies that recognize α-syn_32-46_ peptide from blood at 28 DPI. In both WT and HLA mouse strains, CFA/α-syn_32-46_ immunization produced high levels of antigen-specific antibodies (**Figure S6I**). Thus, CFA/α-syn_32-46_ immunizations in DRB1*15:01 HLA mice trigger an immune response consistent with bulk RNAseq data; however, the α-syn_32-46_ – specific T cells have likely left the peripheral lymphoid organs and are redistributed to other organs, potentially the ENS.

### CD4^+^ T_H_1 / T_H_17 cells adopt a tissue resident memory gene signature in the ileum of CFA/α-syn_32-46_ immunized HLA mice typically found in inflamed mucosal barriers

To determine which immune cell types are responsible for the upregulated pathways identified by bulk RNAseq of the ileum in CFA/α-syn_32-46_, -immunized HLA mice, in particular the adaptive immune response, we performed scRNAseq and analysis on fluorescence-activated cell sorted (FACS) intestinal CD45^+^ immune cells after dissociation of the small intestine from sick CFA/α-syn_32-46_, CFA-immunized, or PBS-injected HLA mice at 21 DPI (**Figure 5A; Table S2A**). We performed normalization, feature selection, and non-linear dimensional reduction of the raw gene expression data after removal of ribosomal (*Rps*/*l*-) and mitochondrial (*Mt*-) genes associated with technical artifacts from enzymatic dissociation. After quality control and filtering, 5430 cells from PBS-injected, 9826 cells from CFA, and 12642 cells from CFA/α-syn_32-46_ -immunized HLA mice were used for downstream analysis (**Table S2A**). We used Harmony batch correction to integrate scRNAseq data from three batches (Korsunsky et al., 2019), and identified 27 unique cell clusters from the three conditions using the non-linear dimensional reduction technique UMAP (**Figures 5D, S6A**). The UMAP analysis revealed large populations of lamina propria T cell subtypes, intraepithelial lymphocytes (IELs) and plasma cells, and smaller populations of myeloid cells, innate lymphoid cells, and memory B cells isolated from the gut in each condition (**Figure 5D, E; Table S2B**).

Since CFA/α-syn_32-46_ immunization in HLA mice triggered constipation and enteric neuronal loss, we postulated that α-syn-specific CD4^+^ T cells may home to the gut, as they were not present in lymphoid tissue (**Figure S5**). To examine transcriptome changes in CD4^+^ T cell subtypes after α-syn_32-46_ immunization, we separated CD4^+^ T_H_1 and T_H_17 cell clusters from the CD45^+^ single cell dataset and performed differential gene expression analysis among three conditions **(Figure 5F, H, K)**. T_H_1, and to a lesser extent T_H_17, cells isolated from CFA/α-syn_32-46_-immunized HLA mice upregulated a gene signature reminiscent of CD4^+^ tissue resident memory cells (T_RM_; high levels for *Crem*, *Vps37b*, *Ramp3*, *Ifngr1* and *Tnfrsf4*; **Figure 5G, I, J, L, M)**; this signature has been identified recently in the murine colon and the murine lung following viral infection (Andreatta et al., 2022; Miragaia et al., 2019; Swarnalekha et al., 2021; Tanemoto et al., 2022). T_H_1 cells from CFA/α-syn_32-46_ immunization also upregulated *Fosl2* expression **(Figure 5J)**, a gene essential for acquisition of an T_RM_ identity [reviewed in (Yenyuwadee et al., 2022)]. Both T_H_1/T_H_17 populations upregulated pro-survival memory genes [e.g., *Bcl2* and *Il7r* (**Figure 5J, M**)]; however, *Id3*, a marker of CD4^+^ memory T_H_1 cells (Shaw et al., 2022) was absent from the T_H_1 population in CFA/α-syn_32-46_-immunized HLA mice, suggesting that they were not typical memory cells. We re-clustered T_H_1 and T_H_17 cells to identify putative subpopulations with distinct T_RM_ signatures **(Figure 5H, K)**. There was an enrichment for T_RM_ gene signatures in subcluster 2 of T_H_1 population, which was over-represented by cells from the CFA/α-syn_32-46_-immunized HLA mice **(Figure 5H, I; Table S2B)**. In contrast, the T_RM_ gene signature was more dispersed among the five T_H_17 subclusters **(Figure 5K, L)**. The T_H_1/T_H_17 cell effector genes and cytokines (*Cd69*, *Ifng* and *Il22*) were also significantly downregulated in T_H_17 cells from CFA/α-syn_32-46_-immunized mice **(Figure 5M),** whereas additional T_RM_ genes such as *Rgs2* (Amezcua Vesely et al., 2019; Li et al., 2016) and *Samsn1* (Gaublomme et al., 2015; Miragaia et al., 2019) were upregulated in T_H_17 cells from CFA/α-syn_32-46_ HLA mice **(Figure 5M)**. Gene-set enrichment analysis (GSEA) of T_H_1 and T_H_17 cells did not reveal significant GO terms associated with genes enriched from CFA/α-syn_32-46_-immunization in HLA mice. Overall, these results demonstrate that CFA/α-syn_32-46_ immunizations in HLA mice triggered transcriptome shifts in CD4^+^ T_H_1 / T_H_17 cells consistent with a tissue resident memory phenotype, similar to the signature found in antigen-experienced T_RM_ cells at mucosal barriers following infection or chronic inflammation. The transcriptome identity shift provides an explanation of why T_H_1 / T_H_17 cells were not found in peripheral lymphoid organs in CFA/α-syn_32-46_-immunized HLA mice (**Figure S5**).

Two populations of IELs (IEL 1, induced IEL) were predominately derived from the CFA/α-syn_32-46_ immunized HLA mice (>45% of the cluster), even after correcting for cell numbers from other conditions **(Figure 5E; S6C, D, F; Table S2B).** These two clusters were characterized by a high expression of effector T cell markers and cytokines (*Itgae*, *Cd69, Gzma, Gzmb, Id2*), but low expression of memory genes (*Id3*, *Tcf7*, *Xcl1*) (**Figure S6F**). The third population of IELs (IEL 2) expressed high levels of CD8^+^ T cell memory identity (*Id3*, *Tcf7*, *Xcl1*), but lacked expression of effector cytokines (*Ifng*, *Gzma*, and *Gzmb*) (**Figure S6F**). In contrast to the CD8α^+^ IEL 1 cluster, the CD8αβ^+^ induced IEL cluster is likely derived from antigen-experienced CD8^+^ T cells which home to intestinal epithelium following activation [reviewed in (Cheroutre et al., 2011)]. Differential gene expression analysis of induced IELs among three conditions revealed upregulation of T_RM_-associated genes (e.g., *Nr4a1*, *Rgs2*, and *Vps37b*) in CFA/α-syn_32-46_ immunized HLA mice (**Figure S6E, F**). Based on this unique gene signature, the induced IELs are likely an effector-like antigen-experienced CD8αβ^+^ IEL population (Wang et al., 2023) highly enriched in CFA/α-syn_32-46_ immunized HLA mice. Granulocytes, composed of CCR3^+^ eosinophils and CXCR2^+^ neutrophils, were predominantly derived from CFA-immunized HLA mice **(Figure S6G).** Differential gene expression analysis revealed upregulated genes associated with granules (*Lcn2*, *Chil3*) and the type 1 interferon response (*Ifitm1*, *Ifitm2*, *Ifitm3*, *Ifitm6*, *Irf1*, *Ifnar2*) in CFA/α-syn_32-46_-compared with CFA-immunized HLA mice **(Figure S6J, K)**. GSEA confirmed enrichment of genes for: a) defense response to bacterium (GO: 0042742); b) response to interferon beta (GO: 0035456); and c) humoral immune response (GO: 0006959) **(Figure S6I)**. Overall, the scRNAseq analysis of immune cells shows that the immune response in the gut at peak constipation (21 DPI; **Figure 5A-C**) was driven not only by CD4^+^ T cells, but also CD8αβ^+^ effector IEL cells and granulocytes, that likely contribute to disease outcomes.

### CD4^+^ T cells are partially responsible for the ENS dopaminergic neuron damage

Since CD4^+^ T cells and CD8αβ^+^ effector IEL cells drive immune responses in the gut of CFA/α-syn_32-46_ immunized HLA mice, we depleted either CD4^+^ or CD8^+^ T cells with anti-CD4, anti-CD8, or isotype control antibodies administered before and at weekly intervals after α-syn_32-46_ immunization to determine which cells are critical for inflammation, neurodegeneration and constipation (**Figure 6A**). Flow cytometry analysis of peripheral blood showed that the appropriate antibody treatments depleted CD4^+^ and CD8^+^ T cell types in mice (**Figure 6B**).

**Figure 6.**
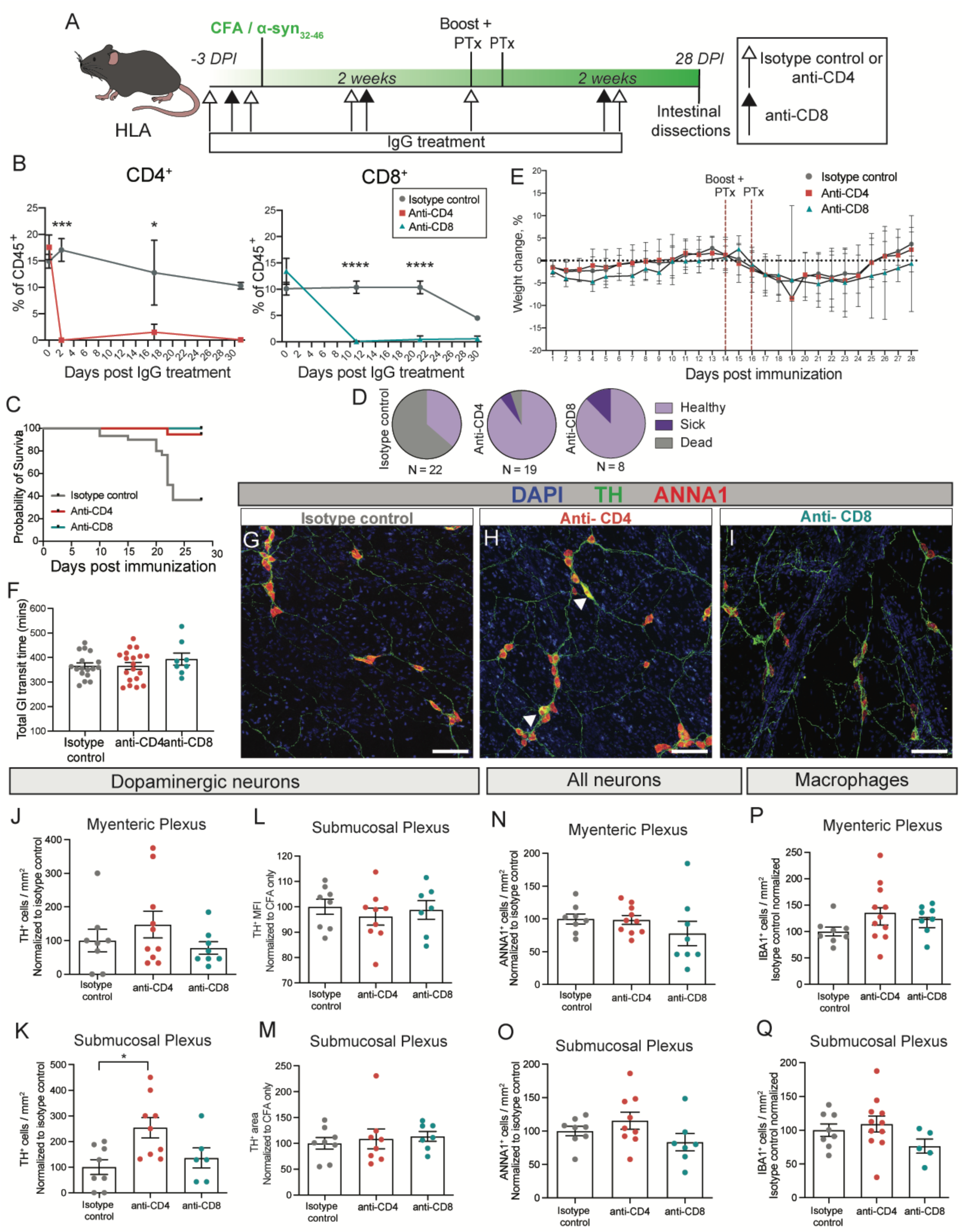
CD4^+^ T cell depletion partially rescues neuronal loss in α-syn_32-46_ immunized HLA mice. **(A)** A schematic diagram of the experimental design. HLA mice were immunized with CFA/α-syn_32-46_ peptide and received a boost after two weeks. At 14 and 16 DPI, *B. pertussis* toxin (Ptx) was administered intravenously. Mice received α-CD4 or isotype control antibodies intraperitoneally at −3 and −1 DPI (white arrows) and then at weekly intervals. Mice received α-CD8 antibodies intraperitoneally at −2 DPI (black arrows) and then at 10-day intervals. At 28 DPI, the ileum was dissected out for analysis. **(B)** Graph showing the levels of CD4^+^ or CD8^+^ T cells as % of CD45^+^ cells at three distinct DPIs in CFA/α-syn_32-46_ HLA mice for each treatment: α-CD4 (red), α-CD8 (teal), or isotype control (gray). **(C)** Kaplan Meier curve depicting the probability of survival and **(D)** pie charts displaying the proportion of α-CD4 (red)-, α-CD8 (teal)-, or isotype control (gray)-treated CFA/α-syn_32-46_ HLA mice that remained healthy (light purple), became ill (dark purple) or died (gray). **(E)** Graph depicting the percent of weight change from the initial weight (0 DPI) of α-CD4 (red)-, α-CD8 (teal)-, or isotype control (gray)-treated CFA/α-syn_32-46_-immunized HLA mice. **(F)** Dotted bar graph depicting the total GI transit time of α-CD4 (red)-, α-CD8 (teal)-, or isotype control (gray)-treated CFA/α-syn_32-46_ HLA mice. **(G-I)** Representative fluorescent images of the submucosal plexus of Isotype control **(G)**, α-CD4 **(H)**, α-CD8 **(I)** treated CFA/α-syn_32-46_ HLA mice stained with DAPI (blue), TH (green) and ANNA1 (red). White arrowheads indicate TH^+^ cell bodies. Scale bar = 100 µm. **(J-Q)** Dotted bar graphs depicting the number of dopaminergic neurons **(J, K)**, TH^+^ MFI **(L)**, TH^+^ area **(M)**, all neurons **(N, O)**, and macrophages **(P, Q)** in the myenteric and submucosal plexuses of α-CD4 (red)-, α-CD8 (teal)-, or isotype control (gray)-treated CFA/α-syn_32-46_ HLA mice. Data analyzed by repeated-measure ANOVA (**B, C**), and two-way ANOVA (**G-N**). Bar graphs depict the mean and error bars the SEM. Each symbol in the bar graphs represents the data collected from one mouse. *p<0.05, ***p<0.001, **** p<0.0001. Isotype control n = 8 mice, α-CD4 n = 10 mice, α-CD8 n = 8, data collected from 3 independent experiments.

A high proportion (∼64%) of CFA/α-syn_32-46_ immunized HLA mice treated with the IgG isotype control did not survive to 28 DPI, while nearly all CD4^+^- or CD8^+^-depleted CFA/α-syn_32-46_- immunized HLA mice survived until sacrifice (**Figure 6C, D**). Weight monitoring revealed no significant difference in weight loss across all treatment groups (**Figure 6E**). All groups showed similar GI transit times (**Figure 6F**) and there was no significant difference in macrophages in either myenteric or submucosal plexuses (**Figure 6P, Q**). Importantly, histopathological analysis revealed a rescue of TH^+^ neurons in the submucosal plexus of CD4^+^-depleted CFA/α-syn_32-46_ HLA mice, but no rescue by the depletion of CD8^+^ T cells (**Figure 6G-K**). Interestingly the neuronal loss appeared to be specific to TH^+^ cells, as other ENS neurons remained unchanged across all treatment groups. Thus, CD4^+^ T cells may be in part responsible for inducing TH^+^ cell damage and loss after CFA/α-syn_32-46_-immunizations in HLA mice, but additional factors like CD8αβ^+^ effector IEL cells, or granulocytes may contribute to other disease phenotypes such as constipation and weight loss.

## DISCUSSION

PD is a multifactorial disease and dysregulation of the immune response targeting α-syn has been strongly implicated as a potential driver of pathogenesis, at least in a subset of PD patients during the first decade after diagnosis (Garretti et al., 2019; Lindestam Arlehamn et al., 2020; Sulzer et al., 2017). Our previous studies have shown that CD4^+^ T cells that respond prominently to two regions of the α-syn protein are found in the blood of PD patients (Sulzer et al., 2017). Although the immune response to the α-syn_32-46_ region is restricted to patients carrying a specific HLA haplotype [B*07:02 C*07:02 DRB5*01 DRB1*15:01 DQA1*01:02 DQB1*06:02] (Sulzer et al., 2017), how these interactions trigger PD pathogenesis is unknown.

Using an active immunization approach in mice similar to induction of the EAE mouse model for human MS (Lengfeld et al., 2017; Lutz et al., 2017), we find that α-syn_32-46_ peptide immunizations in mice in which their native MHC class II allele was substituted by the human HLA DRB1*15:01 allele recapitulate multiple aspects of the immune response detected in PD patients. These include loss of ENS neurons, a progressive decrease in dopaminergic neuronal processes, weight loss and constipation, features reminiscent of clinical gastroenterological deficits reported in PD patients twenty years prior to disease onset (Khoo et al., 2013) (**Figure 7**). Some of these disease phenotypes (e.g., progressive decrease in dopaminergic neurons) require, at least in part, CD4^+^ T cells (**Figure 6**), although the mechanisms by which CD4^+^ T cells mediate them are unclear. Our findings confirm previous hypotheses that α-syn_32-46_ antigen / HLA-DRB1*15:01 interactions underlie some of the prodromal stages of PD manifested by prominent gut inflammation and CD4^+^ T cell-driven loss of enteric neurons.

**Figure 7.**
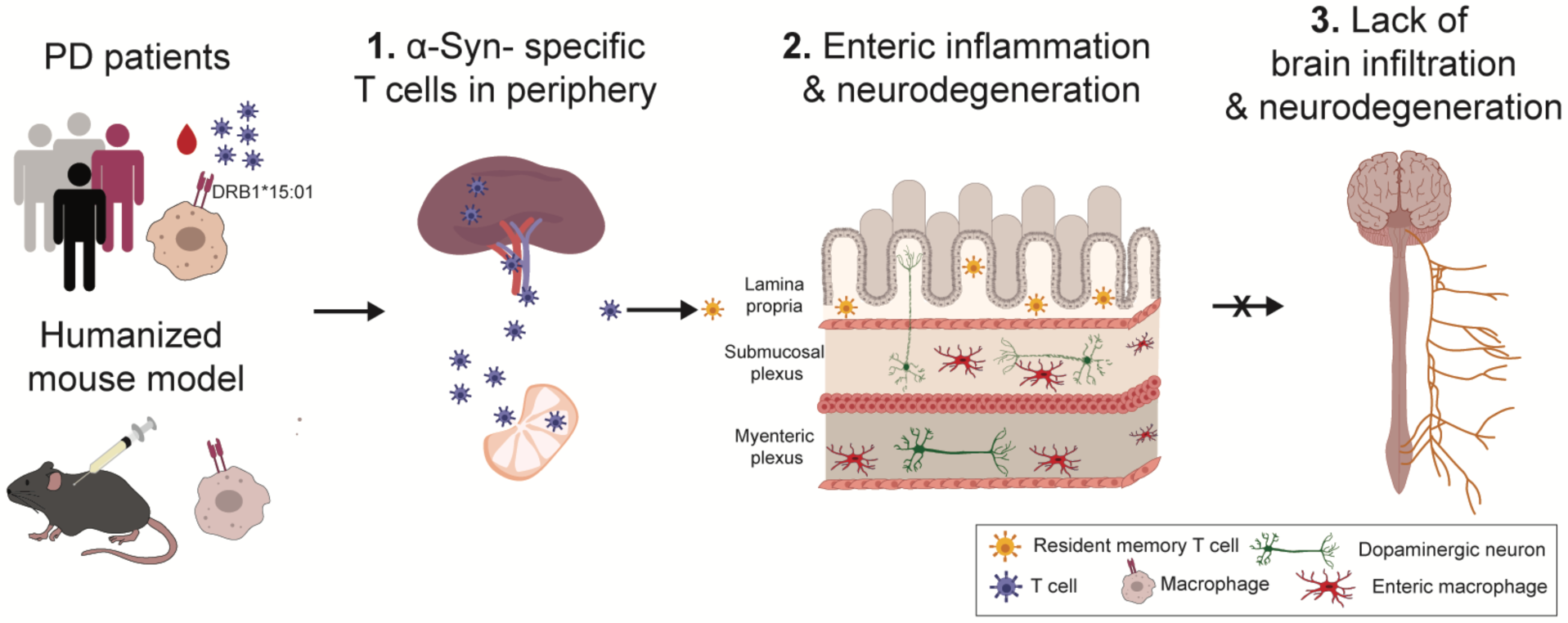
The role of α-syn-specific T cells in PD pathogenesis. α-Syn-specific T cells have been identified in PD patients carriers of the HLA DRB1*15:01 allele. Humanized mice, lacking MHCII^-/-^ and expressing HLA DRB1*15:01, that are immunized with the α-syn_32-46_ peptide may produce α-syn-specific T cells (T_H_1 and T_H_17 lymphocytes; blue) in the peripheral lymphoid organs (spleen and lymph nodes). These T cells (blue) leave the circulation to home to target organs, where they may acquire a putative “resident memory” phenotype (yellow). We postulate that the gut is one such organ where T_H_1/ T_H_17 cells (blue) are converted to mucosal T_RM_ (yellow) found normally during infection or chronic inflammation based on our single cell RNA sequencing data. In addition, signatures of both innate and adaptive immune responses are upregulated in the gut with CFA/α-syn_32-46_ peptide immunizations based on bulk RNA sequencing. The activation of both innate and adaptive immune responses in CFA/α-syn_32-46_-immunized HLA mice may trigger enteric neurodegeneration in the submucosal plexus. However, there is neither inflammation nor T cell infiltration into the brains of α-syn_32-46_-immunized HLA mice, suggesting that additional factors are required to induce the CNS neuropathology and behavioral phenotypes characteristic of PD.

While the immunizations generated a strong antibody response to α-syn_32-46_ in both WT and HLA mice, we found that T_H_1, T_H_17 and CD8^+^ T cells that responded to the peptide were retained in spleen and axillary lymph nodes only in WT mice. Strikingly, α-syn_32-46_ - specific CD4^+^ T cells were absent in immune organs in CFA/α-syn_32-46_ - immunized HLA mice (**Figure S5**). This observation is consistent with effective homing of these CD4^+^ T cells from dedicated immune organs to the gut (McLachlan and Jenkins, 2007), where they are likely responsible for gastrointestinal inflammation, loss of ENS neurons and constipation (**Figure 7**). Single cell RNA sequencing of intestinal immune cells confirmed the presence of CD4^+^ T_H_1 and T_H_17 lymphocytes in the gut of CFA/α-syn_32-46_ peptide immunized mice, where these cells adopted a transcriptome signature characteristic of antigen-experienced tissue resident memory cells found in mucosal barriers during infection and chronic inflammation, including IBD [reviewed in (Lange et al., 2022; Lyu et al., 2022)] (**Figure 5G-M**).

In our humanized mouse model, constipation, a GI symptom present in the majority of PD patients, specifically required the combination of α-syn_32-46_ and the HLA DRB1*15:01 allele. While other PD mouse models have reported constipation (Metzger and Emborg, 2019; Sampson et al., 2016), our study demonstrates that a specific combination of an α-syn-derived epitope and an HLA allele can drive gut pathology and dysmotility. This pathology is characterized by activation of innate and adaptive immune gene signatures (**Figure 5B, C**), which is followed initially by a loss of ANNA1^+^ enteric neurons in the submucosal plexus at 28DPI, and subsequently by a progressive reduction in TH^+^ neuronal processes and intensity between 28-42 DPI in CFA/α-syn_32-46_ HLA mice (**Figures 3, 4, 7**).

Depletion experiments of CD4^+^ and CD8^+^ T cell suggest that CD4^+^ T cells are critical drivers of dopaminergic neuron damage and loss in the gut, and are consistent with other studies implicating CD4^+^ T cells in neuronal damage in other PD mouse models (Brochard et al., 2009; Williams et al., 2021). The timing of the enteric features in this model are consistent with a robust interferon response gene signature revealed by bulk RNAseq (**Figure 5B, C**). While we find that CD4^+^ T cells promote enteric neuron loss, the relationship between CD4^+^ T cell-mediated neuronal loss and interferon response requires further investigation. The interferon response at 21 DPI may reflect dying enteric neurons with an increase in extracellular nucleic acids. Type 1 interferon response have been suggested to play a non-classical role in neurodegeneration, including in the MPTP mouse model for Parkinson’s disease (Main et al., 2016).

Our single cell RNA sequencing has identified a population of CD4^+^ T_RM_ cells present in the gut of sick CFA/α-syn_32-46_-immunized HLA mice at 21DPI. In human IBD, CD4^+^ CD69^+^ CD103^+^ T_RM_ cells are proposed to accumulate and regulate intestinal inflammation (Lyu et al., 2022; Zundler et al., 2019). A similar CD4^+^ T_RM_ population (CD161^+^ CCR5^+^) produces proinflammatory cytokines in Crohn’s disease (Yokoi et al., 2023). Populations of CD103^-^ CD4^+^ T_RM_ cells similar to those identified in our scRNAseq results have been shown to expand following antigen exposure. For example, *Aspergilus fumigatus* generates CD103^-^ CD4^+^ T_RM_ cells in the murine lung, which produce proinflammatory cytokines and contribute to fibrosis [reviewed in (Hirahara et al., 2021)]. In addition to CD4^+^ T_RM_ cells, CD8^+^ CD103^-^ T_RM_ cells in the intestine give rise to CD8^+^ CD103^+^ T_RM_ cells that respond to secondary infections (Fung et al., 2022; von Hoesslin et al., 2022). The identification of CD4^+^ T_RM_ cells in the gut of CFA/α-syn_32-46_-immunized HLA mice at 21 DPI, coupled with the critical role that these cells play in chronic inflammation, suggest that they may play a critical role to drive ENS pathology in the gut.

The overrepresentation of induced IELs and granulocytes from CFA/α-syn_32-46_ -immunized HLA mice suggests that the expansion from antigen exposure may not be limited to CD4^+^ T cells. These additional immune cells may play important roles in the gut pathology following CFA/α-syn_32-46_ immunizations, although this is difficult to ascertain with our small sample size. Other immune cell types are likely involved in the constipation phenotype, and future studies with enrichment of specific immune subtypes could provide additional insights into their role in disease pathogenesis. Granulocytes isolated from CFA/α-syn_32-46_-immunized mice displayed an anti-bacterial phenotype compared to the CFA-immunized mice, since the inducer of anti-bacterial inflammation lipocalin-2 (*Lcn2*) was highly upregulated in this population. *Lcn2* is highly expressed in intestinal inflammation, including IBD [Reviewed in (Moschen et al., 2017)]. Such robust antimicrobial responses in our immunization model may suggest the presence of a leaky gut, which could also contribute to the constipation phenotype. The interferon responses that were highly upregulated from bulk RNA sequencing are largely absent from the single cell dataset, with the exception of granulocytes from CFA/α-syn_32-46_-immunized mice **(Figure S6I, K; Table S2)**. It is likely that other cells in the intestine, such as epithelial cells, may produce the majority of the transcriptional responses detected by bulk RNA sequencing of the ileum. These collective bulk and single cell RNA sequencing data support the model that HLA-α-syn_32-46_ interactions drive innate and adaptive immune responses in the gut that underlie the pathology, and α-syn-specific CD4^+^ T cells are critical, at least in part, to drive the enteric features of prodromal PD.

The mechanisms by which CD4^+^ T cells mediate enteric neuronal loss remain unclear, although their roles have been confirmed in Parkinson’s rodent models with prominent CNS damage (Harms et al., 2021; Williams et al., 2021). The immune-dependent loss of dopaminergic neurites may be transient, and could be related to a temporary loss of the TH^+^ marker expression (Harvey et al., 2000; Hotchkiss and Gibb, 1980; Soriano et al., 1997), as has been shown in other peripheral neurons (Coulombe and Bronner-Fraser, 1986; Wolinsky and Patterson, 1983), although it has also been reported that newborn dopamine neurons may appear in the adult small intestine in response to their death (Kulkarni et al., 2017).

While constipation is strongly associated with prodromal PD, there is little analysis of the inflammatory state or neurodegeneration associated with that condition. The density of enteric neurons in the small intestine is reported to correlate with clinical features of dysmotility (Boschetti et al., 2019). Although dopamine has been shown to be essential for normal gastrointestinal motility in rodents (Li et al., 2006), some biopsy studies of the gut report no difference in dopaminergic neurons between PD patients and healthy controls (Annerino et al., 2012; Corbille et al., 2014; Lebouvier et al., 2010), while others report changes in dopamine in inflammatory bowel disease (Kurnik-Lucka et al., 2021). These analyses are of the colon, and enteric neurons are irregularly distributed across the two plexuses of the 25 feet-long human adult bowel (Rao and Gershon, 2016), making it challenging to thoroughly analyze such specimens. Our study examines up to 1 mm^2^ of myenteric and submucosal plexus of the ileum per mouse at confocal resolution, offering a broad analysis of this enteric region. Importantly, our mouse model examines gut responses to two acute immunizations with a self-antigen, in contrast to chronic antigen exposure present in PD; it is possible that ENS symptoms in PD, including constipation, might wax and wane depending on the presence of specific epitopes, particularly α-syn_32-46_ or phosphorylated S129.

A transient CNS entry of peripheral T cells that recognize a mitochondria-derived antigen associated with intestinal inflammation was reported to drive SN dopamine axonal loss in *Pink1*-deficient mice (Matheoud et al., 2019). In contrast, we observed no CNS symptoms (CD3^+^ T cell infiltrates, neuroinflammation or loss of dopaminergic neurons) in the α-syn_32-46_ / HLA DRB1*15:01 mouse model, consistent with reports that ENS symptoms appear decades prior to CNS symptoms in PD patients. The lack of CNS damage in the α-syn_32-46_ / HLA DRB1*15:01 model suggests that a “second hit” is required to target α-syn-specific T cells into the CNS, followed by neuroinflammation and loss of central dopaminergic neurons. This could be due to the transient, rather than chronic, exposure to the α-syn epitope (two rounds of immunizations in mice in contrast to chronic exposure over decades in PD patients), or insufficient HLA-associated presentation of the α-syn peptide by antigen presenting cells in the brain, which may in turn result from disease-associated altered degradation pathways that produce neoepitopes.

In conclusion, we have shown that α-syn autoimmunity is sufficient to induce constipation and gut pathology with loss of enteric neurons that can elicit prodromal symptoms of PD. The demonstration that a restrictive HLA allele in combination with an α-syn-derived antigen leads to prodromal PD-like symptoms suggests that autoimmune responses to α-syn-derived epitopes may play an early role in PD pathogenesis.

## ACKNOWLEDGEMENTS

We thank Michael D. Gershon, Alcmene Chalazonitis-Green, and Wanda Setlik (CUIMC) for guidance with enteric dissections, staining, and behavior experiments, and particularly with the analysis of neuronal number and area in the flat mount ileum preparations; Ilir Agalliu (AECOM) for his advice with advanced statistical analysis; Peter Sims (CUIMC) for his guidance with the analysis of scRNAseq data; Ori Lieberman, Michael Post, Charlotte Wayne, Madeline Edwards, Megan Sykes, Ai Yamamoto, and Wassim Elyaman (CUIMC) for suggestions with various experiments throughout this study. This research was funded in part by Aligning Science Across Parkinson’s [#0375] through the Michael J. Fox Foundation for Parkinson’s Research (MJFF). For the purpose of open access, the authors have applied a CC BY public copyright license to all Author Accepted Manuscripts arising from this submission. D.S and E.K are also supported by NIH/NINDS (R01 NS095435) and the JPB Foundation. T.C., S.S., and D.A., are also supported by grants from the NIH/NIMH (R01 MH112849), NIH/NIE (R01 EY033994), the National MS Society (RG-1901-33218). D.A., and T.C., are partially supported by unrestricted gifts from the Newport Equities LLC, the Walz family and PANDAS Network to the Department of Neurology, CUIMC.

## AUTHOR CONTRIBUTIONS

F.G., D.A., A.S., and D.S., conceived the project; F.G., C.M., N.S., S.S., S.W.K., T.C., E.K. and D.A. performed the experiments and analyses, F.G., C.M., T.C. A.S., D.A., and D.S. wrote and edited the manuscript, and F.G. and C.M. prepared figures and figure legends.

## STAR METHODS

### RESOURCE AVAILABILITY

#### Lead Contact

Further information and requests for resources and reagents should be directed to and will be fulfilled by the lead contact, David Sulzer.

#### Materials availability

This study did not generate new unique reagents.

#### Data and code availability

##### Data

Microscopy reported in this paper will be shared by the lead contact upon request. RNA-sequencing data will be made available on the NCBI Gene Expression Omnibus (GEO) database.

##### Code

This paper does not report original code.

##### Section 3

Any additional information required to reanalyze the data reported in this paper is available from the lead contact upon request.

### EXPERIMENTAL MODEL AND SUBJECT DETAILS

#### Mice

All experimental procedures were approved by the Institutional Animal Care and Use Committee (IACUC) at Columbia University Irving Medical Center (CUIMC). Generation of transgenic mice expressing the *HLA DRB1*15:01* α and β chains in the C57BL/6 background have been previously described (Krogman et al., 2017). Wild-type C57BL/6J were obtained from the Jackson Laboratory (Bar Harbor, ME). The HLA transgenic mice lacking mouse MHC-II were backcrossed to the C57BL/6J strain (Jackson Laboratory; Bar Harbor, ME) for more than 8 generations. All mice were bred within the barrier facility of CUIMC and group housed, fed regular chow. Both female and male mice were used in this study in equal numbers at the ages described in the text.

### METHOD DETAILS

#### Immunizations

8- to 12-week-old male and female mice were immunized subcutaneously with a 100 μl emulsion of 100 μg α-syn_32-46_ peptide (A&A peptides; San Diego, CA) dissolved in PBS with Complete Freund Adjuvant (CFA) containing 200 μg of *M. tuberculosis* H37Ra (BD Difco Adjuvant, Fisher Scientific, Cat #DF0638-60-7; Waltham, MA). The day of immunization was designated as day 0. Mice received a boost with the same immunization dose two weeks later. Mice received intravenous injections of 400 ng of *B. pertussis* toxin (List Biological Laboratories, Cat #181; Campbell, CA) 14- and 16-days post-initial immunization (day 0 and 2 post boost). Control animals received *B. pertussis* toxin and emulsion containing PBS/CFA without the protein or the α-syn peptide. Additional control mice received subcutaneous injections of PBS only. Mice were monitored for their weight, food and fluid intake and neurological deficits and their weights were recorded daily. Mice were classified as sick if they displayed 12% reduction from 0 DPI weight and/or hunched posture with ruffled and ungroomed fur.

#### Gut dissections

Mice were perfused with PBS and segments of the ileum were collected in PBS. The contents were flushed out with PBS and the preparations were cut open along the mesenteric border. The flat tissue was stretched tautly and pinned flat on Sylgard (Sylgard^TM^ 184, Dow Corning, Cat #240-4019862; Midland, MI) with the mucosa surface facing up. Specimens were fixed for one hour at room temperature 4% paraformaldehyde and washed three times with PBS for a total of 30 min. The mucosa and submucosal plexus were removed from specimen by microdissection. Then, the mucosal was separated from the submucosal layer. The longitudinal muscles were dissected from the specimen to reveal the myenteric plexus. The submucosal and myenteric plexuses were collected and store in PBS at 4°C until immunofluorescence staining.

#### Intestinal flat mount immunofluorescence staining and analysis

##### Staining

For immunofluorescence analysis, intestinal flat mounts were blocked and permeabilized for 1 hour at room temperature with 10% normal donkey serum (Jackson Immunoresearch, Cat #017-000-121; West Grove, PA) and 1% Triton-X in PBS. Intestinal flat mounts were then incubated overnight at room temperature with primary antibodies in 10% normal donkey serum, 1% Triton-X in PBS. Sections were stained with antibodies against tyrosine hydroxylase (TH) (1:500, Millipore-Sigma, Cat # AB152; Burlington, MA) and CD3 (1:400, Bio-Rad Laboratories, Cat #MCA2690; Hercules, CA), ANNA1 (1:32,000; kind gift by the Gershon laboratory (Margolis et al., 2016)), Iba1 (1:500, WAKO, Cat # 019-19741; Richmond, VA), CD68 (1:1000, Abcam, Cat # ab53444; Waltham, MA). Secondary antibodies with the appropriate conjugated fluorophores were purchased from Invitrogen. Images were acquired with an LSM700 confocal microscope and processed with Fiji software (ImageJ, NIH; Bethesda, MD).

##### Acquisition

For enteric neuron imaging, both the myenteric (MP) and submucosal plexus (SP) for each mouse were analyzed. Within each plexus of each animal, 2-3, 2×2 tile z-stack 640.17x 640.17 µm confocal images at 20x magnification were acquired. This totaled to over 1 mm^2^ of flat-mount enteric tissue imaged and analyzed per mouse. For imaging macrophages, both the MP and SP were analyzed for each animal. Within each plexus of each animal, 2-3, z-stack 390.09 x 390.09 µm confocal images at 20x magnification were acquired.

##### Analysis

The investigator who analyzed the images was blinded to each condition. The number of ANNA1^+^, TH^+^ cells, and IBA1^+^ cells were counted for each image within each layer. Within the SP, the TH^+^ signal was thresholded and the area and MFI of the thresholded TH^+^ signal was analyzed for mean fluorescent intensity and size of area. The thresholding was consistent across each image, animal, and condition within each experiment. Within each animal, the number of ANNA1^+^, TH^+^ cells, and IBA1^+^ cells across all images were summed and divided by acquisition area. Within each experiment, the cell numbers were normalized to CFA only.

#### Total GI transit

Carmine red, which cannot be absorbed from the lumen of the gut, was used to study total gastrointestinal (GI) transit time (Kimball et al., 2005). A solution of carmine red (300 μl; 6%; Sigma-Aldrich, Cat #C1022; St Louis, MO) suspended in 0.5% methylcellulose (Sigma-Aldrich, Cat #M0512; St Louis, MO) was administered by gavage through a 21-gauge round-tip feeding needle. The time at which gavage took place was recorded as *T*_0_. After gavage, fecal pellets were monitored at 10 min intervals for the presence of carmine red. Total GI transit time was considered as the interval between *T*_0_ and the time of first observance of carmine red in the stool.

#### Bulk RNA sequencing of the ileum

Ileum samples were incubated in TRIzol reagent (Thermo Fisher Scientific, Cat #15596026; Waltham, MA) and stored at −80 _C. Total RNA was extracted and assessed for quality utilizing Agilent bioanalyzer for quantitation of RNA/DNA/protein (CUIMC Molecular Pathology Core). cDNA library preparation and RNA sequencing were performed by the CUIMC Genome Center. mRNAs were enriched from total RNA samples with poly-A pull-down, then processed with library construction using Illumina TruSeq chemistry. Libraries were then sequenced using Illumina NovaSeq6000 at CUIMC Genome Center. Samples were multiplexed in each lane, which yielded targeted number of paired-end 100bp reads for each sample. RTA (Illumina) for base calling and bcl2fastq2 (version 2.19) were used for converting BCL to fastq format, coupled with adaptor trimming. A pseudoalignment was performed to a kallisto index created from transcriptomes (GRCm38) using kallisto (0.44.0). Differentially expressed genes under various conditions were tested using DESeq2R packages designed to test differential expression between two experimental groups from RNA-seq counts data. Genes were considered differentially expressed if they had an adjusted p-value <0.05 and a log_2_fold change below or above 0.5. The differential expression was normalized for each gene.

#### Single cell RNA sequencing of the ileum

##### Isolation of intestinal immune cells

The methods used for isolation of intestinal immune cells were adapted from Ivanov et al. (Ivanov et al., 2006). Briefly, mice were perfused with PBS and the small intestine was removed. Peyer’s patches were dissected from the small intestines, and the intestine was cut open longitudinally. Intestinal contents were removed by scraping along the tissue with rounded forceps, followed by 5-6 washes with cold PBS. The intestine was cut into large fragments, then placed in 5-10 mL of cell dissociation solution (5mM EDTA, 10mM HEPES, in HBSS supplemented with 2.5% FBS) and incubated for 10 minutes at 37°C rotating at 100 rpm. After 10 minutes, the solution was vortexed for 25 seconds, then the supernatant was collected and discarded using a wire mesh to collect the tissue. The tissue was incubated again in fresh cell dissociation solution, then the additional supernatant was discarded as previously described. The tissue fragments were rinsed in HBSS then cut into small pieces (∼1 mm^2^). The small fragments were incubated in three 20-minute intervals at 37 C with slow rotation in 5 mL of digestion solution (0.5 mg/mL collagenase D (Roche, Cat# 11088858001; Mannheim, Germany), 0.5 U/mL Dispase (Stem Cell Technologies, Cat# 07913; Vancouver, Canada), 0.5 mg/mL DNase I (Roche, Cat# 10104159001; Mannheim, Germany), in RPMI supplemented with 5% FBS). After each 20-minute digestion period, the solution was vortexed for 30 seconds, then the supernatant was collected by filtering through a 100 µm strainer. The remaining tissue fragments were placed in new digestion solution for subsequent washes. All supernatants were combined then resuspended with 10 mL of 40% Percoll (Sigma-Aldrich, Cat# GE17-0891-01; St. Louis, MO) in RPMI with 10% FBS. The resuspended cells were gently added on top of 5 mL 80% Percoll in RPMI with 10% FBS in a 15 mL conical tube. The tubes were centrifuged at 2500 rpm for 20 minutes without brake. The cell layer at the interface between 40% and 80% Percoll was collected, diluted in RPMI with 10% FBS. The cells were pelleted, then resuspended in FACS buffer, consisting of 1% bovine serum albumin (Sigma-Aldrich, Cat# A9647-100G; St. Louis, MO) in PBS.

##### FACS

The cells isolated from the small intestine were resuspended with 400 µl Fc block (BD Biosciences, Cat# 553142; Franklin Lakes, NJ), 1:200 in FACS buffer, and incubated for 20 minutes on ice. The cells were washed in 500 µl FACS buffer, pelleted at 1500 rpm for 2 minutes, then resuspended in 600 µl of FACS buffer with 1:100 CD45 BUV395 antibody (BD Biosciences, Cat# 564279; Franklin Lakes, NJ) and incubated on ice for 30 minutes. After 30 minutes, the cells were washed in 500 µl FACS buffer, then resuspended in 500 µl of FACS buffer containing propidium iodide (BioLegend, Cat# 421301; San Diego, CA) at 1:10,000. Viable, CD45^+^ cells were sorted by the Sony MA900 Multi-Application Cell Sorter.

##### Analysis

The isolated CD45^+^ intestinal immune cells were processed by the Single Cell Analysis Core at Columbia University using the TruSeq Stranded mRNA Library Prep Kit (Illumina) and sequenced on an Illumina NovSeq 6000 platform. The Single Cell Analysis Core used 10X Cell Ranger analysis software (version 5.0.1) to process fastq files using “cellranger count” for each sample individually with default parameters producing an alignment of reads to the mm10-2020-A transcriptome. The gene-cell matrices were processed, including normalization, scaling, selection of features, linear dimensional analysis, and clustering, using the Seurat package v4.0.0 in R Studio (Hao et al., 2021). For quality control, cells with fewer than 15% mitochondrial reads and UMI counts between 1000 and 50000 were kept for downstream analysis. Ribosome (*Rps*/*l*-) and mitochondria (*Mt*-) associated genes were regressed from the dataset to remove effects of technical artifacts of the cell dissociation process. Batch correction was performed with the Harmony package (Korsunsky et al., 2019). UMAP clusters were manually annotated by assessment of differentially expressed features using the FindMarkers function in the Seurat package **(Figure S6B; Table S2C)**.

#### T cell depletions

Mice were injected interperitoneally with 250 μg of Ultra-LEAF Purified anti-mouse CD4 or Rat IgG2b isotype control antibodies (Biolegend, Cat #100457 and Cat #400644; San Diego, CA) 3 and 1 days prior to immunization. After immunization, mice were injected with 250 ug of antibody weekly. Cell depletion was assessed via flow cytometry. For depletion of CD8^+^ T cells, mice were injected interperitoneally with 250 μg of Ultra-LEAF Purified CD8^+^ or Rat IgG2b isotype control antibodies (Biolegend Cat # 100746 and Cat #400644; San Diego, CA) 2 days prior to immunization. After immunization, mice were injected with 250 ug of each antibody 9- and 19-days post immunization, then cell depletion was determined via flow cytometry.

## QUANTIFICATION AND STATISTICAL ANALYSIS

Fiji software (ImageJ, NIH; Bethesda, MD) was utilized for the quantitation of immunofluorescence images. Statistical analysis was performed using GraphPad Prism software version 8.0 (Graphpad Software Inc; San Diego, CA). For enteric immunofluorescence analysis, we compared ANNA1^+^ and TH^+^ cell density and TH^+^ area and MFI among 3 immunization groups: PBS, CFA only, or CFA/α-syn_32-46_ using one way analysis of variance (ANOVA). In addition, we used Scheffe method for multiple comparisons given that 3 groups had unequal sample size. We then fitted a general linear regression with immunization group as exposure and adjusted for variability from two different raters (FG vs. CM). All tests were 2-sided with significant level set at p<0.05. For all other analyses, unpaired t test was used to compare the differences between two groups in studies. One-way and two-way ANOVA followed by Bonferroni post hoc tests were performed to compare the differences among groups. Differences between groups were presented as the mean ± SEM or SD as noted in the figure legends. The sample size (n) for each experiment is indicated in figure legends and the Results section. All tests were two sided and p values < 0.05 were considered to be statistically significant.

## KEY RESOURCES TABLE

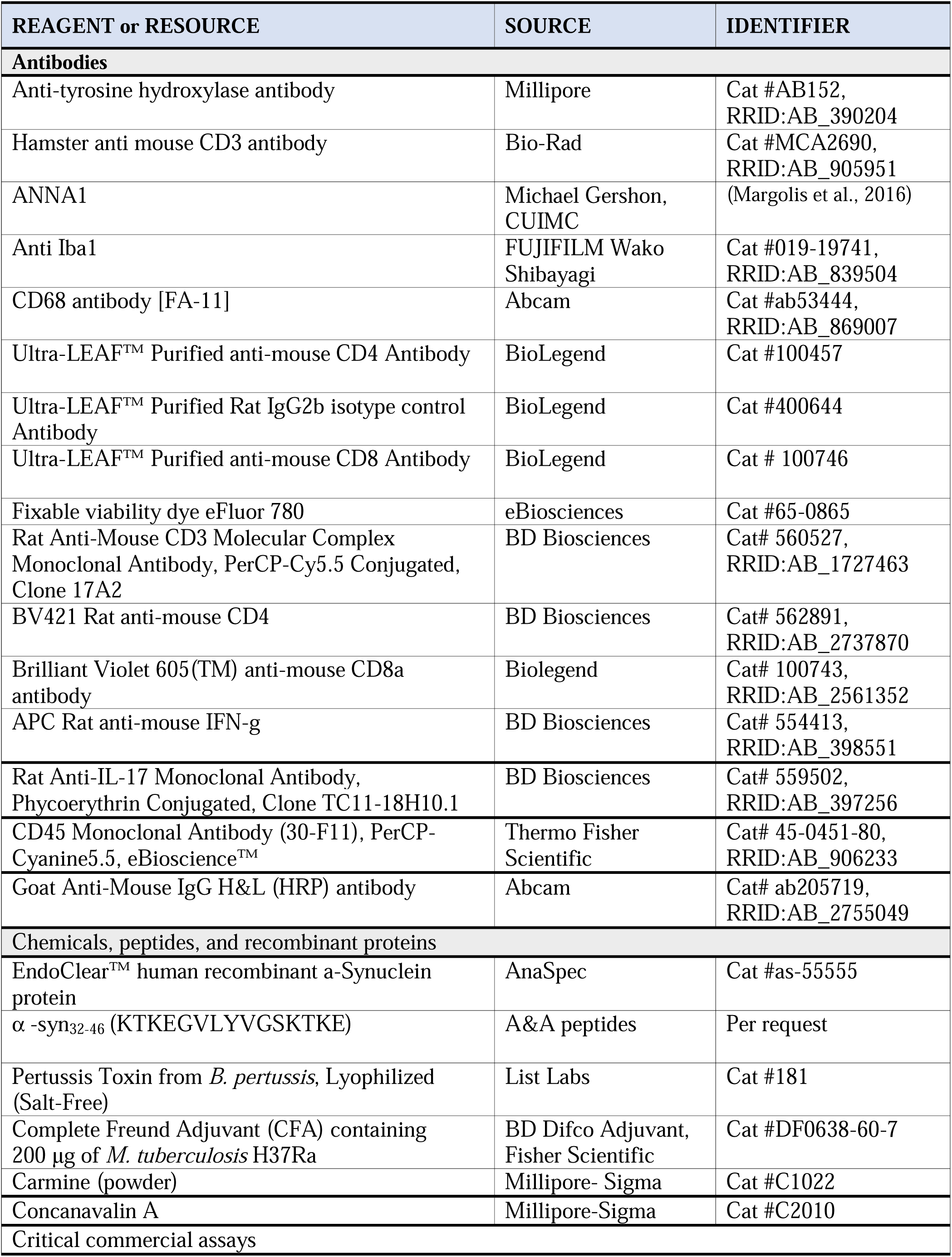
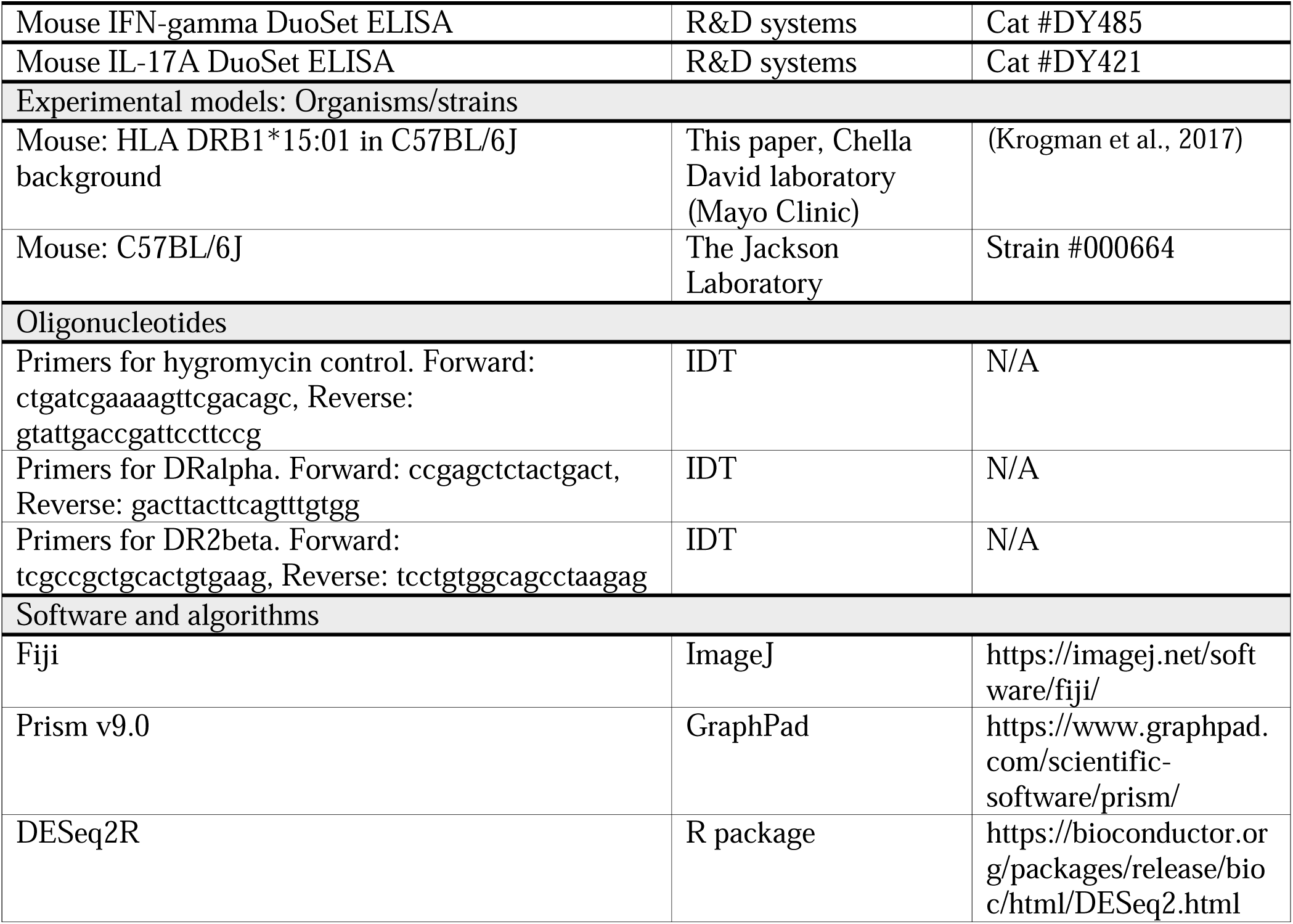

