## Supplemental Data for "Interaction of an α-synuclein epitope with HLA-DRB1*15:01 triggers enteric features in mice reminiscent of prodromal Parkinson’s disease"

### SUPPLEMENTAL FIGURES AND FIGURE LEGENDS

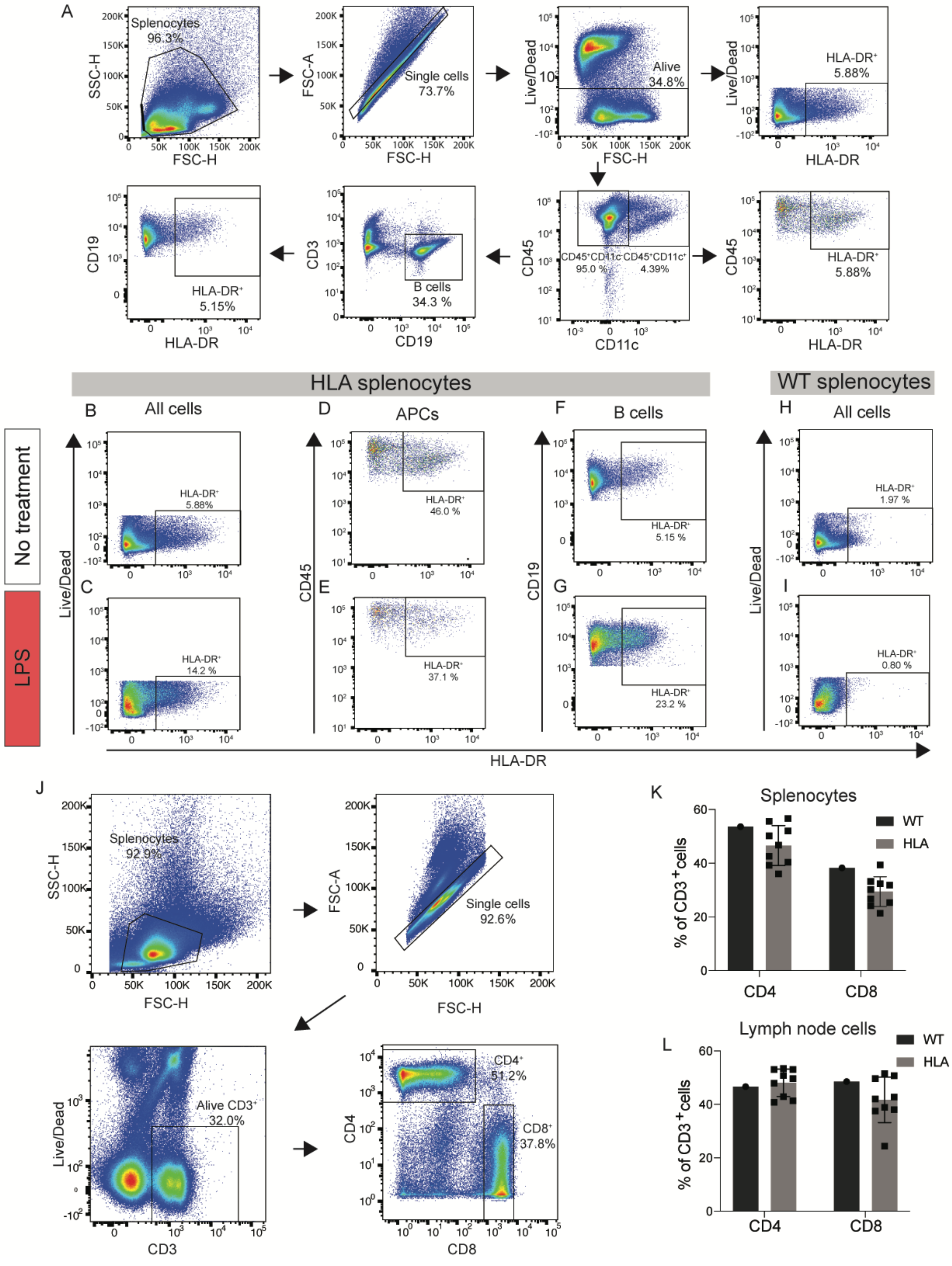

**Figure S1 (linked to Figure 1). Distinct immune cells in the HLA-DRB1*15:01 transgenic mice express HLA-DR, and upregulate its expression upon LPS exposure. (A)** Representative flow cytometry plots show the gating strategy for the HLA-DR protein levels in HLA-DRB1*15:01 mice. After gating for living single cell splenocytes, the HLA-DR protein levels at the membrane were analyzed in all live cells, $CD45^{+}CD11c^{+}$ antigen-presenting cells (APC), and $CD45^{+}CD11c^{-}CD19^{+}$ B cells. (**B-G**) Representative flow plots show HLA-DR expression in live cells **(B, C)**, APCs **(D, E)**, and B cells **(E, F)** isolated from splenocytes of HLA-DRB1*15:01 transgenic mice that were either unstimulated (**B, D, F**), or stimulated with 50 ng/ml lipopolysaccharide (LPS; **C, E, G**). (**H, I**) Representative flow plots show HLA-DR protein surface levels in live cells from the spleen of WT mice that were either unstimulated **(H)** or stimulated with LPS **(I)**. **(J)** Representative flow cytometry plots show the gating strategy for the $CD4^{+}$ and $CD8^{+}$ proportions of $CD3^{+}$ T cells. After gating for single cell splenocytes, alive $CD3^{+}$ T cells were selected, and the percentages of $CD4^{+}$ and $CD8^{+}$ T cells were determined within the $CD3^{+}$ T cell population. **(K)** Dotted bar graph representing the percentages of $CD3^{+}$ T cells that are $CD4^{+}$ and $CD8^{+}$ isolated from either WT (dark bars), or HLA (light bar), splenocytes. **(L)** Dotted bar graph representing the percentages of $CD3^{+}$ T cells that are $CD4^{+}$ and $CD8^{+}$ isolated from either WT (dark bars), or HLA (light bar), lymph node cells. Each dot for the graphs represents one mouse; two-way ANOVA with Sidak multiple comparisons test.

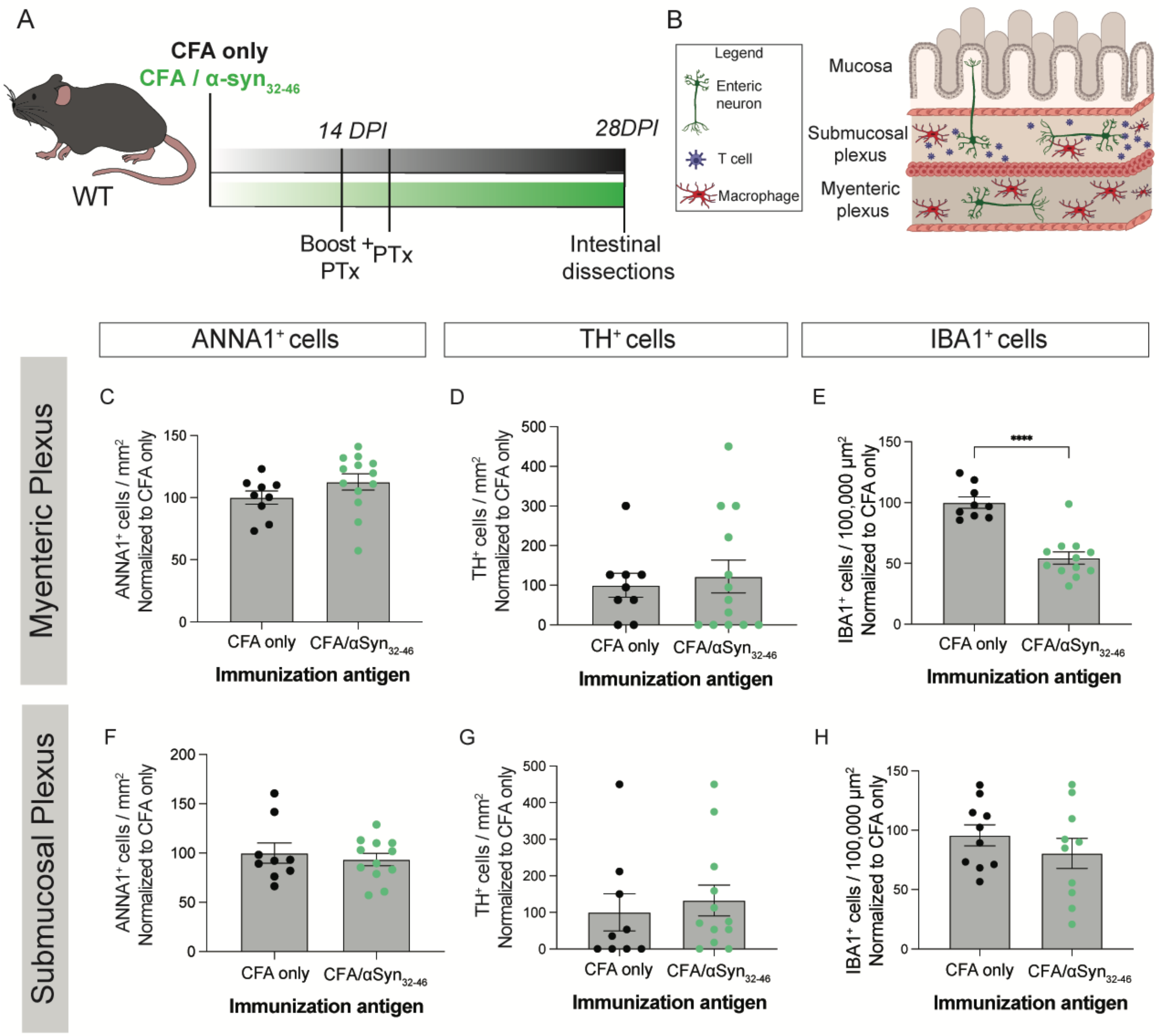


**Figure S2 (linked to Figure 2). α-Syn$_{32\text{-}46}$ immunizations do not induce enteric neuronal loss in wild-type mice. (A, B)** Schematic diagram of the experimental design and the myenteric and submucosal plexuses of the ileum used for the flat mount analysis. **(C-H)** Dotted bar graphs of ANNA1$^+$ and TH$^+$ neurons per mm$^2$, and IBA1$^+$ cells per 100,000 μm$^2$ in the myenteric (**C-E**), and in submucosal (**F-H**) plexuses of CFA only (black circles) and CFA/α-syn$_{32\text{-}46}$ (green circles)-immunized wild-type (WT) mice. Bar graphs show the mean +/- SEM. (**C-H)** Data were analyzed by Mann-Whitney test; **** $p<0.0001$; n=9 CFA WT mice; n=12 CFA/α-syn$_{32\text{-}46}$ WT mice.

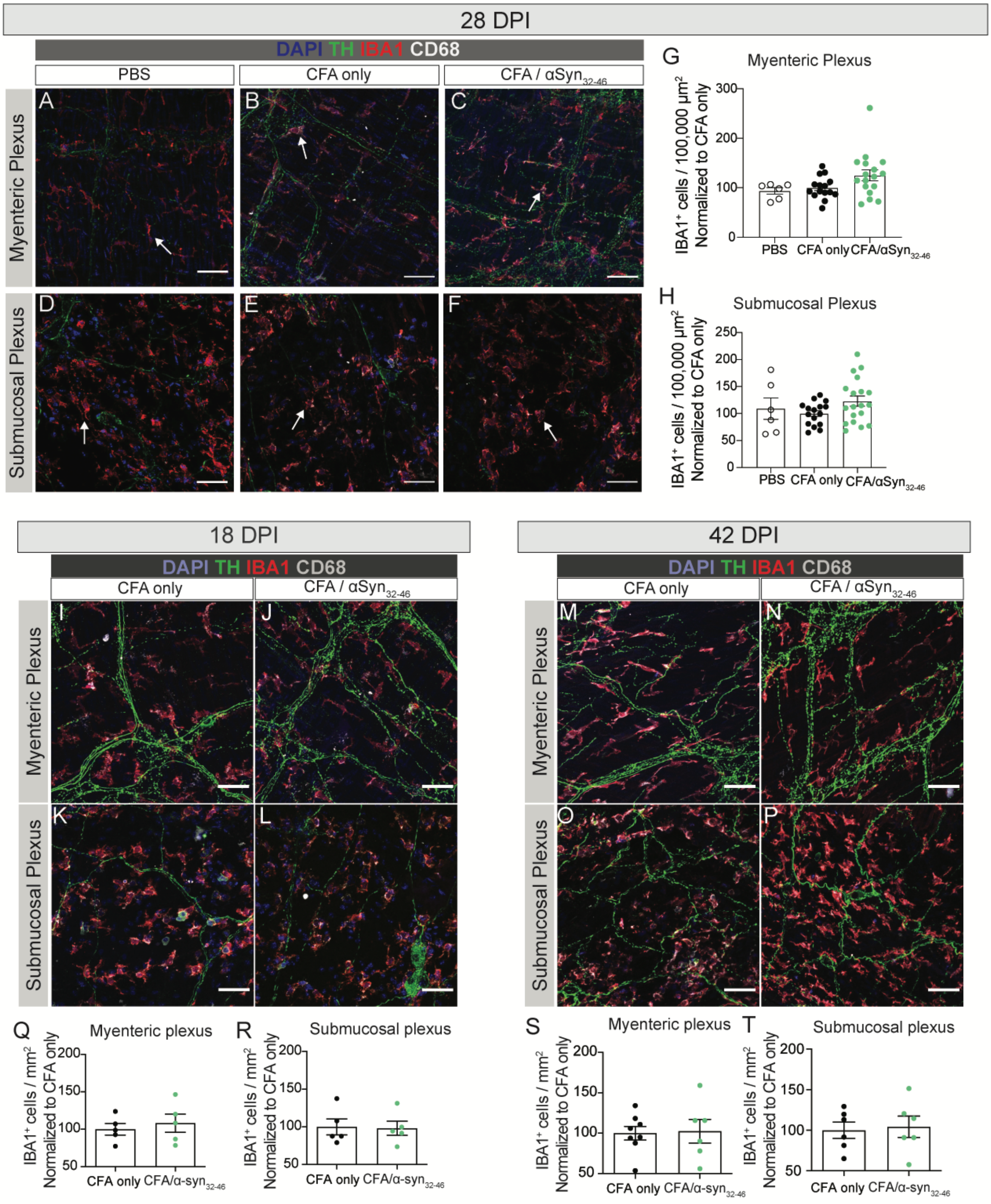


**Figure S3 (linked to Figures 3 and 4). α-Syn32-46 peptide immunizations do not increase macrophage proliferation in the HLA DRB1*15:01 mice. (A-F)** Representative images of the myenteric and submucosal plexuses from three conditions (PBS; CFA only; CFA/α-syn32-46) at 28 days post immunization

(DPI) stained for DAPI (blue), TH (green), Iba1 (red) and CD68 (white). White arrows indicate macrophages. Scale bars = 50 μm. **(G, H)** Dotted bar graphs of the number of $Iba1^+$ cells/100,000 $\mu m^2$ normalized to CFA-only in the myenteric **(G)**, and submucosal **(H)** plexuses of PBS (clear circles), CFA only (black circles), and CFA/α-$syn_{32-46}$ (green circles)-immunized HLA mice. Data were analyzed by Mann-Whitney test; n=6 PBS HLA mice; n=16 CFA only HLA mice; n=19 CFA/α-$syn_{32-46}$ HLA mice. There was no significant difference among the three groups. **(I-L)** Representative images of the myenteric and submucosal plexuses from two conditions (CFA only and CFA/α-$syn_{32-46}$) at 18 days DPI stained for DAPI (blue), TH (green), Iba1 (red) and CD68 (white). Scale bars = 50 μm. **(Q, R)** Dotted bar graphs of the number of $Iba1^+$ cells/100,000 $\mu m^2$ normalized to CFA only in the myenteric **(Q)**, and submucosal **(R)** plexuses of CFA only (black circles) and CFA/α-$syn_{32-46}$ (green circles)-immunized HLA mice. Data were analyzed by Students' t-test; n=5 CFA only HLA mice; n=5 CFA/α-$syn_{32-46}$ HLA mice. There was no significant difference between the two groups. **(M-P)** Representative images of the myenteric and submucosal plexuses from two conditions (CFA only and CFA/α-$syn_{32-46}$) at 42 DPI stained for DAPI (blue), TH (green), Iba1 (red) and CD68 (white). Scale bars = 50 μm. **(S, T)** Dotted bar graphs of the number of $Iba1^+$ cells/100,000 $\mu m^2$ normalized to CFA only in myenteric **(S),** and submucosal **(T)** plexuses of CFA only (black circles) and CFA/α-$syn_{32-46}$ (green circles)-immunized HLA mice. Data were analyzed by Students' t-test; n=8 CFA only HLA mice; n=6 CFA/α-$syn_{32-46}$ HLA mice. There was no significant difference between the two groups.

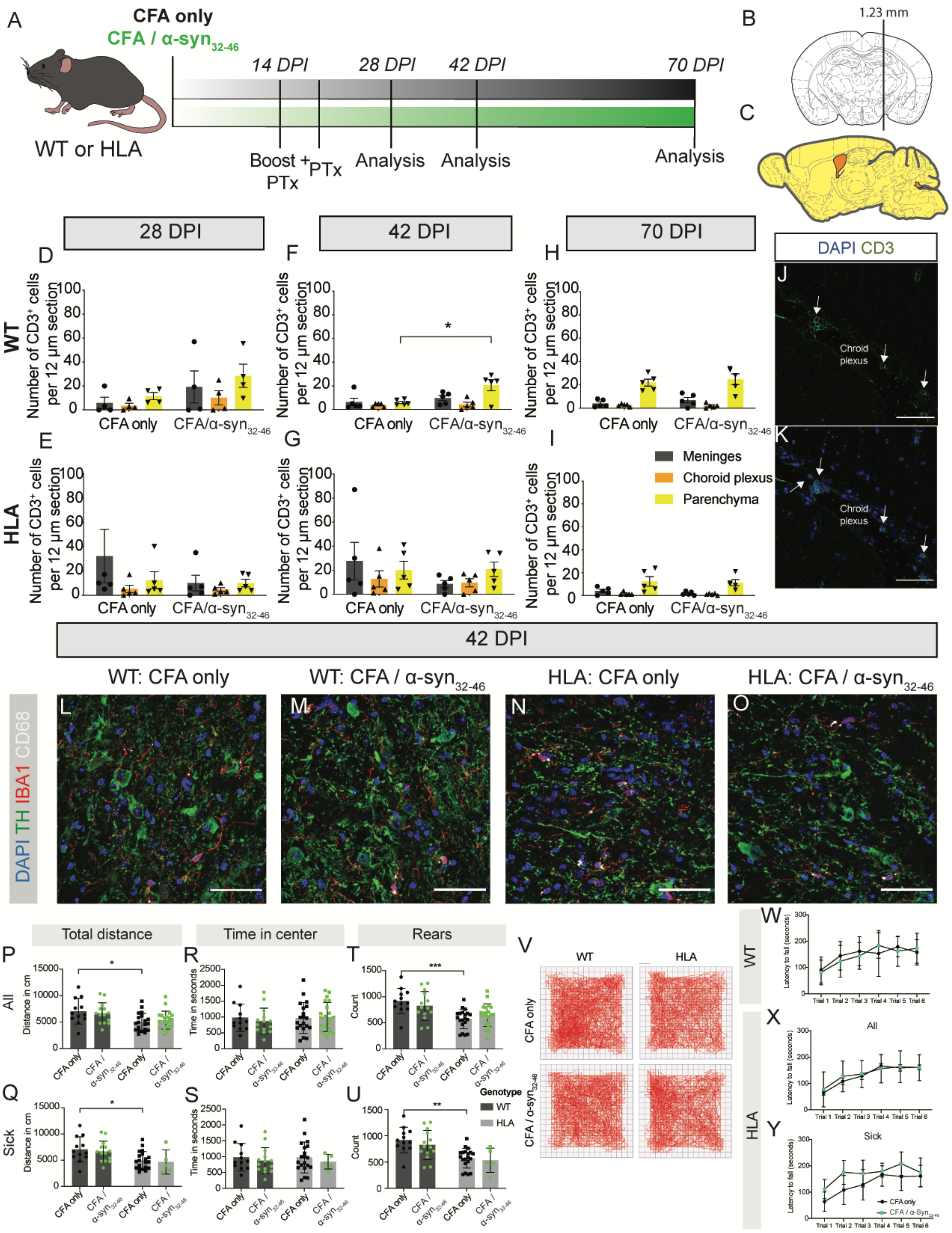
A
CFA only
CFA / α-syn32-46
WT or HLA
14 DPI
28 DPI
42 DPI
70 DPI
Boost + PTx
PTx
Analysis
Analysis
Analysis
B
1.23 mm
C
28 DPI
42 DPI
70 DPI
WT
HLA
Number of CD3+ cells per 12 µm section
CFA only
CFA/α-syn32-46
Meninges
Choroid plexus
Parenchyma
DAPI CD3
Chroid plexus
42 DPI
WT: CFA only
WT: CFA / α-syn32-46
HLA: CFA only
HLA: CFA / α-syn32-46
DAPI TH IBA1 CD68
Total distance
Time in center
Rears
All
Sick
Distance in cm
Time in seconds
Count
Genotype
WT
HLA
CFA only
CFA / α-syn32-46
Latency to fall (seconds)
Trial 1 Trial 2 Trial 3 Trial 4 Trial 5 Trial 6

**Figure S4 (linked to Figure 4). T cells do not infiltrate into the brain parenchyma in CFA/α-syn$_{32-46}$-immunized wild-type or HLA-DRB1*15:01 mice up to 70 days post immunization. (A)** A schematic diagram of the experimental design. Wild-type (WT) and HLA mice were immunized with either CFA only or CFA/α-syn$_{32-46}$ emulsion and received a boost after two weeks. At 14 and 16 DPI, mice were administered an intravenous injection of *B. pertussis* toxin (Ptx) to transiently open the blood-brain barrier. Mice were sacrificed at 28, 42 and 70 days post-immunization (DPI) and brains were collected for analysis. **(B, C)** Brains were sectioned sagittally past +1.23 mm bregma region to include the substantia nigra (SN) region. (**D-I**) CD3$^{+}$ T cells were quantified in each section and their location [meninges (grey), parenchyma (yellow), and choroid plexus (orange)] was noted. Dotted bar graphs of CD3$^{+}$ T cell number in WT (**D, F, H**) and HLA (**E, G, I**) brains located in either meninges (grey), parenchyma (yellow), or choroid plexus (orange) at 28 **(D, E)**, 42 **(F, G)**, and 70 **(H, I)** DPI. **(J, K)** Representative images of CD3$^{+}$ T cell infiltrates (white arrows) in the choroid plexus. (**L-O**) Representative images of the substantia nigra of mice 42 DPI stained for DAPI (blue), TH (green), IBA1 (red), and CD68 (white). (**P-U**) Bar graphs depicting the total traveled distance **(P, Q)**, the time in center **(R, S),** and the total number of rears **(T, U)** in CFA only (black symbols) and CFA/α-syn$_{32-46}$ (green symbols)-immunized WT (dark grey) and HLA (light grey) mice within 60 minutes in the open field test chamber. All mice are included in top panels **(P, R, T)** and only sick, CFA/α-syn$_{32-46}$ -immunized HLA mice are included in bottom panels **(Q, S, U). (V)** Representative traces of the open field test. (**W-Y**) Graphs depicting the latency to fall in seconds in six trials runs across two days for WT **(W)**, all HLA **(X)** and sick only HLA **(Y)** mice. Data were analyzed by either two-way ANOVA or repeated measures ANOVA followed by Bonferroni post-hoc correction; * $p<0.05$, ** $p<0.01$, *** $p<0.001$. Data **D-O**: 28 DPI: WT: CFA only n=4 mice, CFA/α-syn$_{32-46}$ n=4 mice, HLA: CFA only n=4 mice, CFA/α-syn$_{32-46}$ n=4 mice; 42 DPI : WT: CFA only n=5 mice, CFA/α-syn$_{32-46}$ n=5 mice, HLA: CFA only n=5 mice, CFA/α-syn$_{32-46}$ n=5 mice; 72 DPI : WT: CFA only n=5 mice, CFA/α-syn$_{32-46}$ n=5 mice, HLA: CFA only n=5 mice, CFA/α-syn$_{32-46}$ n=5 mice. Bar graphs show mean and error bars the SEM. Data

**P-Y**: WT: CFA only n=12 mice, CFA/α-$syn_{32-46}$ n=14 mice. HLA: CFA only n=20 mice, CFA/α-$syn_{32-46}$ n=21 mice. WT: 3 independent experiments; HLA: 4 independent experiments. Each circle represents the data collected from one mouse. Dotted bar graphs display mean +/- SEM.

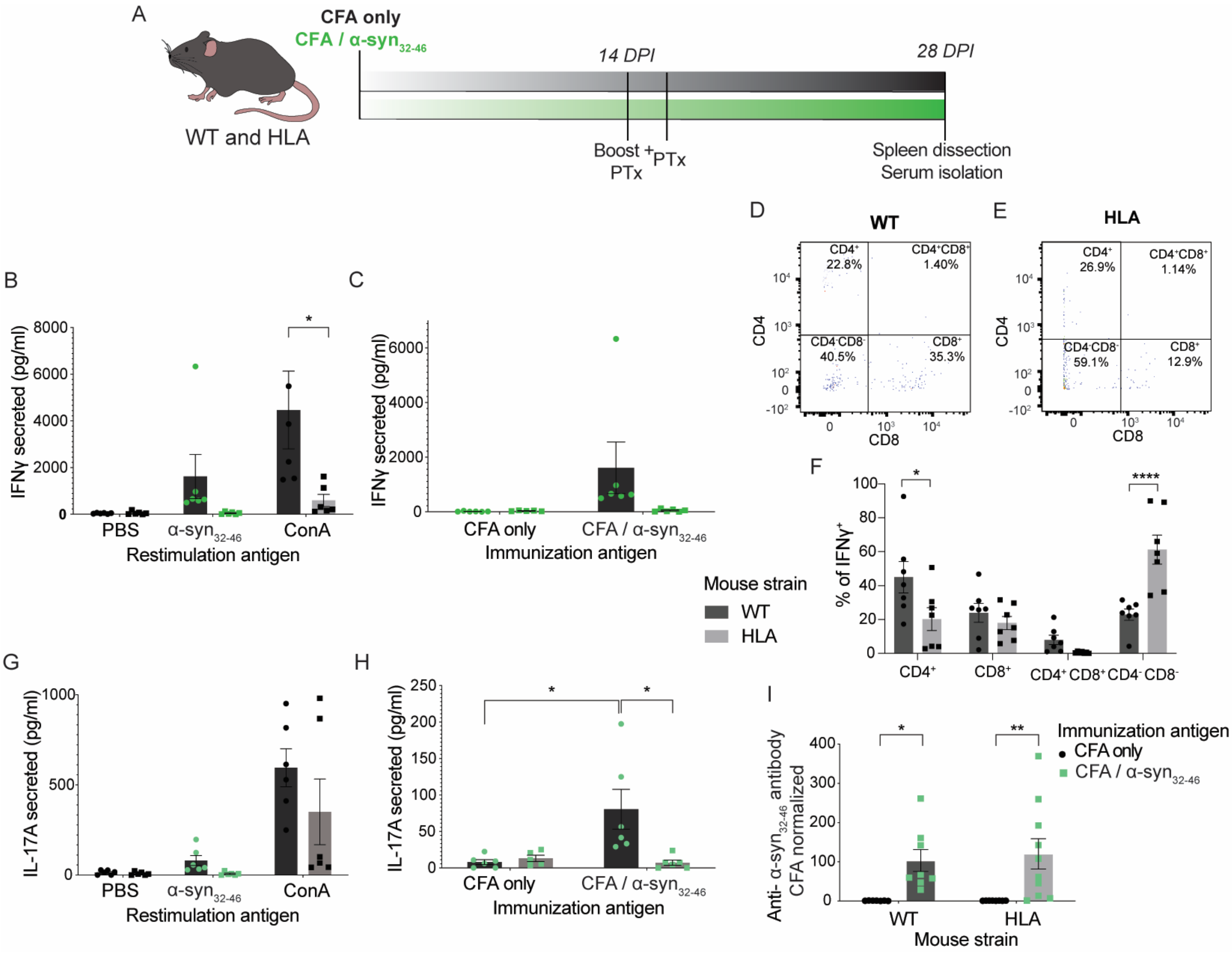


**Figure S5 (Linked to Figure 6). α-Syn$_{32-46}$-specific T cells are found in the circulation in wild-type, but not HLA-DRB1*15:01, mice. (A)** A schematic diagram of the experimental design. Wild-type (WT) and HLA DRB1*15:01 mice were immunized with either CFA only or CFA/α-syn$_{32-46}$ emulsion and received a boost after two weeks. At 14 and 16 DPI, an intravenous injection of *B. pertussis* toxin (Ptx) was administered. At 28 DPI, mice were sacrificed and the spleens collected for *in vitro* re-stimulation assays. Cells were re-stimulated with the immunization antigen (α-syn$_{32-46}$), PBS and ConA as negative and positive controls, respectively, for two days. After two days *in vitro*, the media was collected for IFNγ detection via ELISA **(B, C)** and the cells were collected for ICS analysis via flow cytometry **(D-F)**. **(B)**

Dotted bar graph of IFNγ levels secreted by splenocytes from either CFA/α-$syn_{32-46}$ WT (dark bars), or HLA (light bars) mice after re-stimulation *in vitro* with PBS, α-$syn_{32-46}$ peptide (green) and ConA. **(C)** Dotted bar graphs of IFNγ levels secreted by splenocytes from CFA only and CFA/α-$syn_{32-46}$-immunized WT (dark bars) and HLA (light bars) mice after re-stimulation *in vitro* with the α-$syn_{32-46}$ peptide (green symbols). FACS plots (**D, E**) and dotted bar graph (**F**) of distinct subsets of $CD3^{+}$IFNγ-producing splenocytes that were $CD4^{+}$, $CD8^{+}$, $CD4^{+}CD8^{+}$, or $CD4^{-}CD8^{-}$ as determined by flow cytometry in WT **(D)**, and HLA **(E)** mice. **(G)** Dotted bar graph of IL-17A levels secreted by splenocytes from α-$syn_{32-46}$ -immunized WT (dark bars) and HLA (light bars) mice after re-stimulation *in vitro* with PBS, α-$syn_{32-46}$ peptide (green symbols) and ConA. **(H)** Dotted bar graphs of IL-17A levels secreted by splenocytes from CFA only and CFA/α-$syn_{32-46}$ WT (dark bars) and HLA (light bars) mice after re-stimulation with α-$syn_{32-46}$ peptide (green symbols). **(I)** Dotted bar graph depicting anti-α-$syn_{32-46}$ antibody levels, detected via indirect ELISA, in the sera of CFA only (black circles) and CFA/α-$syn_{32-46}$ (green squares) immunized WT and HLA mice. Data were analyzed by two-way ANOVA followed by post-hoc Bonferroni correction; *$p<0.05$, **$p<0.01$. Splenocyte re-stimulation experiments (**B-H**): WT: CFA only n= 7 mice, CFA/α-$syn_{32-46}$ n=7 mice, HLA: CFA only n=6 mice, CFA/α-$syn_{32-46}$ n=7 mice. WT: 7 independent experiments; HLA: 6 independent experiments. Serum experiments (**i**): WT: CFA only n= 8 mice, CFA/α-$syn_{32-46}$ n=8 mice, HLA: CFA only n=8 mice, CFA/α-$syn_{32-46}$ n=10 mice. WT: 3 independent experiments; HLA: 4 independent experiments. Each dot represents data collected from one mouse. Bar graphs display mean +/- SEM.

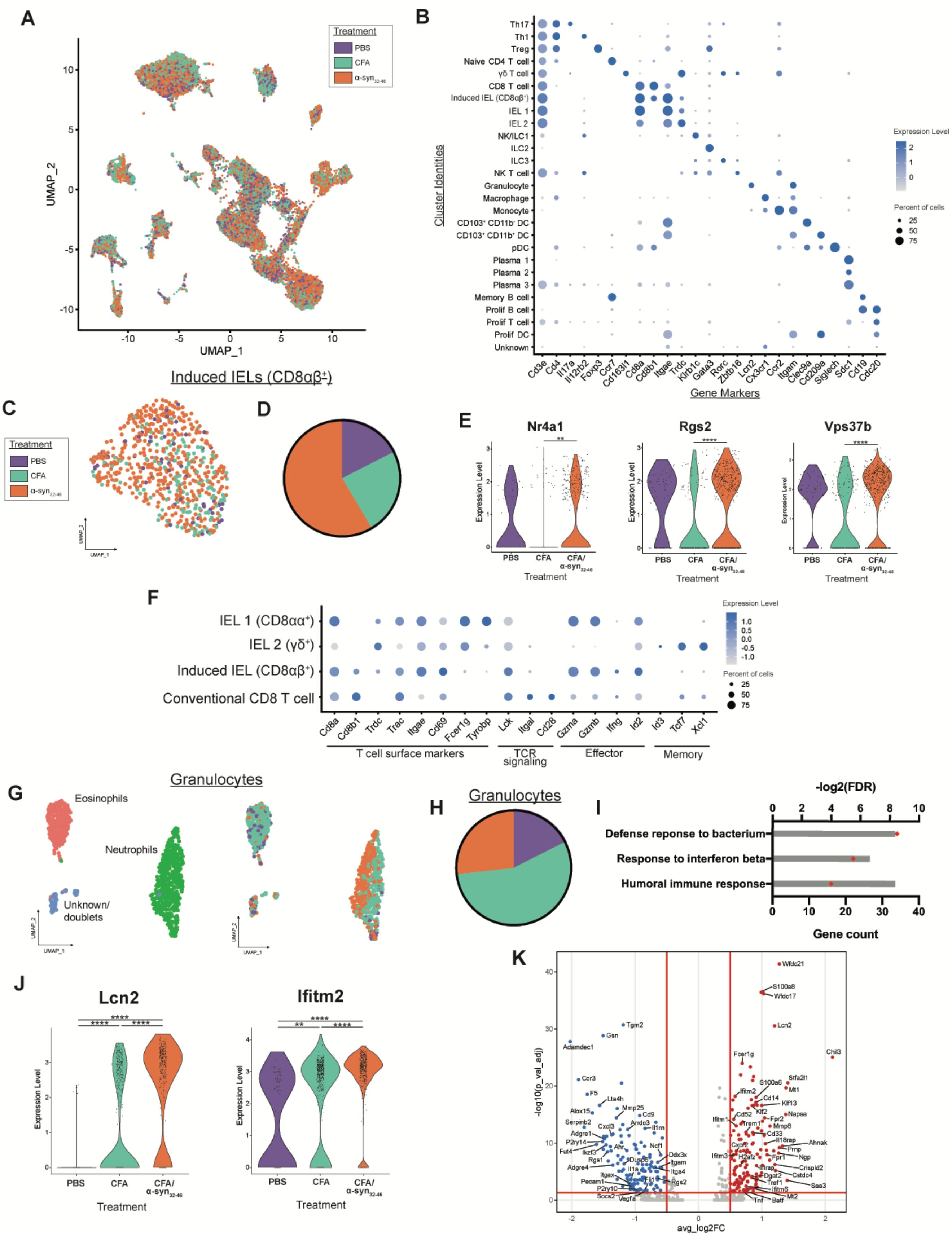
A
Treatment
PBS
CFA
α-syn32-46
UMAP_2
UMAP_1
B
Th17
Th1
Treg
Naive CD4 T cell
γδ T cell
CD8 T cell
Induced IEL (CD8αβ+)
IEL 1
IEL 2
NK/ILC1
ILC2
ILC3
NK T cell
Granulocyte
Macrophage
Monocyte
CD103+ CD11b- DC
CD103+ CD11b+ DC
pDC
Plasma 1
Plasma 2
Plasma 3
Memory B cell
Prolif B cell
Prolif T cell
Prolif DC
Unknown
Cluster Identities
Cd3e Cd4 Il17a Il12rb2 Foxp3 Ccr7 Cd163l1 Cd8a Cd8b1 Itgae Trdc Klrb1c Gata3 Rorc Zbtb16 Lcn2 Cx3cr1 Ccr2 Itgam Clec9a Cd209a Siglech Sdc1 Cd19 Cdc20
Gene Markers
Expression Level
Percent of cells
Induced IELs (CD8αβ+)
C
D
E
Nr4a1
Rgs2
Vps37b
Expression Level
PBS
CFA
CFA/ α-syn32-46
Treatment
F
IEL 1 (CD8αα+)
IEL 2 (γδ+)
Induced IEL (CD8αβ+)
Conventional CD8 T cell
Cd8a Cd8b1 Trdc Trac Itgae Cd69 Fcer1g Tyrobp Lck Itgal Cd28 Gzma Gzmb Ifng Id2 Id3 Tcf7 Xcl1
T cell surface markers
TCR signaling
Effector
Memory
Granulocytes
G
Eosinophils
Neutrophils
Unknown/ doublets
H
Granulocytes
I
-log2(FDR)
Defense reponse to bacterium
Response to interferon beta
Humoral immune response
Gene count
J
Lcn2
Ifitm2
Expression Level
PBS
CFA
CFA/ α-syn32-46
Treatment
K
-log10(p_val_adj)
avg_log2FC
Wfdc21
S100a8
Wfdc17
Lcn2
Chil3
Tgm2
Gsn
Adamdec1
Ccr3
F5
Lta4h
Alox15
Mmp25
Cd9
Serpinb2
Cxcl3
Arrdc3
Il1rn
Adgre1
P2ry14
Ahr
Ncf1
Fut4
Ikzf3
Rgs1
Dusp6
Ddx3x
Itgam
Itga4
Rgs2
Adgre4
Itgax
Il1a
Pecam1
P2ry10
Socs2
Vegfa
Fcer1g
S100a6
Stfa2l1
Mt1
Ifitm2
Cd14
Klf13
Cd52
Klf2
Napsa
Ifitm1
Trem1
Fpr2
Mmp8
Cd33
Il18rap
Ahnak
Cxcr2
Prnp
Ifitm3
H2afz
Fpr1
Ngp
Il1rap
Crispld2
Dgat2
Cstdc4
Traf1
Ifitm6
Saa3
Tnf
Batf
Mt2

**Figure S6 (linked to Figure 5). Intestinal intraepithelial lymphocytes and granulocytes show a unique transcriptome signature in CFA/α-syn$_{32\text{-}46}$ -immunized HLA mice. (A)** UMAP plot of single cell RNA sequencing data from sorted CD45$^{+}$ cells in CFA/α-syn$_{32\text{-}46}$-immunized (orange), CFA only-immunized (teal), and PBS-injected (purple) HLA mice. **(B)** Dot plot showing marker gene (x-axis) expression in each cluster (y-axis) from the UMAP. Darker blue dots indicate higher average expression of that gene in the specified cell type. The size of the dot represents the percent of cells in that group that express the specified marker. **(C)** UMAP plot of isolated induced intraepithelial lymphocytes (IELs; CD8αβ$^{+}$) from CFA/α-syn$_{32\text{-}46}$-immunized (orange), CFA only-immunized (teal), and PBS-injected (purple) HLA mice. **(D)** Pie chart of the relative abundance of induced IEL population within each condition (PBS-purple, CFA-teal, CFA/α-syn$_{32\text{-}46}$-orange). **(E)** Violin plots of *Nr4a1*, *Rgs2*, and *Vps37b* gene expression in the induced IEL population, separated by condition. Statistical analyses were done with the Wilcoxen Rank Sum test, followed by Bonferroni correction for multiple comparisons. **(F)** Dot plot of the CD4$^{-}$ CD8α$^{+}$ lymphocyte clusters and the relative expression T cell surface markers and genes involved in TCR signaling, effector functions, and memory functions. **(G)** UMAP plots of isolated granulocytes, split by sub-cluster identities (left) and conditions (right). **(H)** Relative abundance of granulocyte population within each treatment [CFA/α-syn$_{32\text{-}46}$-immunized (orange), CFA only-immunized (teal), and PBS-injected (purple)]. **(I)** Significant gene ontology terms (GO: 0042742, GO: 0035456, GO: 0006959) identified by gene set enrichment analysis of differentially expressed genes upregulated in CFA/α-syn$_{32\text{-}46}$ compared with CFA-immunized HLA mice. Gray bars indicate -log$_2$(FDR) (upper x-axis), and the red diamonds indicate the number of genes in the GO term (lower x-axis). **(J)** Violin plots of *Lcn2* and *Ifitm2* gene expression in the granulocyte population, separated by condition. Statistical analyses were done with the Wilcoxen Rank Sum test, followed by Bonferroni correction for multiple comparisons. **(K)** Volcano plot of differentially expressed genes in granulocytes from single cell RNA seq data. Genes significantly upregulated (avg_log2FC > 0.5, p_val_adj < 0.05) in CFA/α-syn$_{32\text{-}46}$- compared to CFA-immunized HLA mice are labeled in red, those downregulated (avg_log2FC < -0.5, p_val_adj < 0.05) are labeled in blue. Statistical significance in violin plots: * $p_{adj} < 0.05$, ** $p_{adj} < 0.01$, *** $p_{adj} < 0.001$, **** $p_{adj} < 0.0001$.
